# A PLUM Job: Peptide modeLs for Understanding and engineering antiMicrobial therapeutics

**DOI:** 10.64898/2026.02.21.707214

**Authors:** Priyanka Banerjee, Iddo Friedberg, Britta Rued, Oliver Eulenstein

## Abstract

**Motivation:** Antibiotic-resistant infections in humans and animals are rising, creating an urgent need for new antimicrobial strategies. This challenge extends from human health to food production, where foodborne pathogens cause substantial animal and human illness annually. Antimicrobial peptides (AMPs) are promising alternatives because of their broad activity and potentially lower resistance risk. However, rational AMP design remains challenging, particularly when generating targeted peptides with desired function and length for downstream experimental testing.

**Results:** We introduce *Peptide modeLs for Understanding and engineering antiMicrobial therapeutics* (PLUM), a structured conditional variational autoencoder for controlled AMP generation. PLUM organizes the latent space into components associated with function, length, and residual sequence context, enabling *de novo* and prototype-guided generation across the 5–35 amino acid range. In *de novo* generation, PLUM achieved the highest classifier-confirmed yields for both AMP and non-AMP targets. PLUM-generated AMPs were also substantially distinct from the training set and exhibited broad internal sequence diversity. They further showed favorable predicted safety, stability, and potency profiles, including low toxicity, the lowest median instability index among the evaluated models, and a large number of candidates predicted to have high antibacterial potency. In prototype-guided generation, PLUM yielded more AMP-classified variants than HydrAMP while also supporting non-AMP-targeted generation from existing prototypes. Together, these results support PLUM as a scalable framework for function- and length-controlled AMP design.

**Availability:** https://github.com/priyamayur/PLUM

## Introduction

Antimicrobial peptides (AMPs) typically consist of 10–50 amino acids, serving as the first line of defense against invading pathogens. They are found in all domains of life and have been extensively studied for their broad-spectrum antimicrobial activities (Lei et al., 2019; Zhang and Gallo, 2016). The discoveries of cecropins in 1981 and magainins in 1987 marked some of the earliest breakthroughs in AMP research (Zhang and Gallo, 2016). In recent decades, the widespread overuse of conventional antibiotics and the resulting escalation of antibiotic resistance have renewed interest in AMPs as potential therapeutic alternatives. Due to their broad activity spectrum, thermal stability, and immunological effects, AMPs represent a promising class of candidates for next-generation antimicrobial agents for human and animal health (Zhang et al., 2025).

AMPs have recently garnered interest as a novel method for reducing infections with problematic bacterial species, particularly in livestock and healthcare settings. Alternative methods are environmental controls such as surface decontamination and identification of contaminated sources (Obe et al., 2023; Haque et al., 2020). While these strategies have been successful in reducing the incidence of related outbreaks, they do not eliminate their occurrence and are complicated by the systemic overuse of antibiotics. Therefore, regulators in the United States and the European Union have begun to heavily monitor or have outright banned the application of specific antibiotics in livestock, particularly due to the increase in antimicrobial resistance (Abreu et al., 2023). As a result, the need for alternative strategies and methods to control bacterial infections has increased. Initial research has demonstrated that AMPs are a viable alternative to modulate pathogen colonization and provide protective immunological benefits (Robles Ramirez et al., 2024; Oliveira Júnior et al., 2025). The use of AMPs provides a novel strategy for controlling bacterial infections in humans and animals. The goal of designing *Peptide modeLs for Understanding and engineering antiMicrobial therapeutics* (PLUM) was to enable controllable, large-scale generation of novel, easily synthesized linear AMPs for these applications, representing a departure from the typical use of current models.

Traditionally, the discovery of novel antimicrobial peptides has relied on iterative experimental modification testing and optimization in a laboratory setting. (Yan et al., 2022; Cesaro et al., 2022). The experimental modification approach is often time-consuming and cannot explore a diverse range of peptide sequences (Szymczak et al., 2023; Das et al., 2018). Therefore, there is a opportunity for developing efficient computational strategies that can accelerate AMP discovery and facilitate the generation of novel, diverse AMP candidates.

Generative machine learning models have gained popularity for learning complex data distributions and creating realistic samples in images, language, and audio. This capability has inspired their use in peptide design, enabling AMP generation. Early machine learning research on AMPs primarily focused on sequence classification using traditional algorithms (Xiao et al., 2013; Bhadra et al., 2018; Joseph et al., 2012; Su et al., 2019; Veltri et al., 2018). With deep learning, attention shifted toward generative modeling for designing novel peptides with desired properties. Long short-term memory networks (LSTMs) were among the first models applied to peptide sequence generation (Nagarajan et al., 2018; Muller et al., 2018; Wang et al., 2021), followed by VAEs, generative adversarial networks (GANs), Transformers, and diffusion models, driving a surge in AI-guided peptide design. VAEs and GANs were early generative approaches (Tucs et al., 2020; Ferrell et al., 2020; Szymczak et al., 2023; Das et al., 2018). Namely, PepGAN generated both AMPs and non-AMPs, while AMP-GAN introduced conditional control. Conditional VAEs (cVAEs) enabled property-guided generation (Das et al., 2018; Dean and Walper, 2020; Szymczak et al., 2023); PepCVAE followed standard cVAE frameworks (Das et al., 2018), and HydrAMP improved diversity through latent space disentanglement (Szymczak et al., 2023). More recently, diffusion models have provided highly controllable generation (Wang et al., 2024; Cao et al., 2025), with ProT-Diff leveraging ProtT5-guided diffusion and TG-CDDPM generating AMPs from text prompts (Wang et al., 2024; Cao et al., 2025).

Many existing models have demonstrated success in *de novo* AMP generation. However, most focus primarily on AMP candidate generation and do not jointly support functional modulation, non-AMP generation, length control, and variant generation from known peptides. While AMP generation is important, practical applications—such as exploring diverse peptide space or producing sequences with controllable properties—require models that enable flexible and targeted peptide design. HydrAMP is one of the few models capable of generating both AMPs and non-AMPs and one of the few to explore prototype-conditioned analogue generation. Although it provides a valuable starting point for comprehensive peptide design, HydrAMP conditions sequence generation on *E. coli* minimum inhibitory concentration (MIC), which may bias outputs toward a single species, and is limited to producing peptides up to 25 amino acids. Additionally, these models have largely ignored aspects such as toxicity and protease stability, rendering the designed AMPs unusable in vivo even if they inhibit pathogens in vitro.

PLUM is a structured cVAE model for peptide design, inspired by advances in representation learning (Klys et al., 2018; Chen et al., 2019; Esmaeili et al., 2019; Mathieu et al., 2019; Cheng et al., 2020). While standard cVAEs typically encode multiple attributes within a shared latent space, assigning attribute-associated information to dedicated latent components can support a more interpretable organization of the latent representation and enable more targeted control during generation. PLUM extends this principle to peptide sequences by organizing the latent space into components associated with functional activity, length, and residual sequence context. This structured design allows control of peptide properties during generation while maintaining flexibility and diversity. Building on this architecture, PLUM introduces capabilities that address key limitations of existing models, offering four novel major contributions:

1. Structured latent architecture: PLUM organizes the latent space into components associated with function, length, and residual sequence context supporting targeted manipulation of peptide properties.
2. *De novo* generation across functional classes and lengths: PLUM generates both AMPs and non-AMPs across the 5– 35 amino acid range, enabling large-scale exploration of functionally distinct peptide sequences.
3. Prototype-guided generation: PLUM can generate new peptide variants from a given prototype sequence while controlling target functional activity and length.
4. Antibacterial potency-based prioritization: PLUM includes an antibacterial potency model (APM) that prioritizes generated AMP candidates with stronger predicted antibacterial activity.

Together, these features establish PLUM as a flexible, structured platform for controlled exploration and design of peptide sequences beyond conventional AMP-focused approaches.

## Methods

This section describes the datasets, PLUM generative framework, peptide generation methods, and classifier-based assessment used in this study. We first describe the curated AMP and non-AMP datasets used for model training, then present PLUM as a structured generative model for controlled AMP design. We next outline the two peptide generation modes and the downstream classifiers used to characterize generated peptides. Together, these components define a methodological pipeline for data-driven peptide generation and evaluation.

### Dataset Construction

We curated a set of positive (antimicrobially active) and negative (non-antimicrobially active) peptide sequences to train PLUM. We harvested esiting curated datasets of natural linear AMPs (5–35 residues) and matched non-AMPs, along with associated activity data (MICs). AMP sequences were collected from the AMP databases CAMP, DRAMP, DBAASP, and GRAMPA (Gawde et al., 2023; Ma et al., 2025; Pirtskhalava et al., 2021; Witten and Witten, 2019). We then filtered in the peptides that were linear, containing canonical amino acids only, and sequence validity, yielding 4,723 unique sequences. The non-AMP dataset was retrieved from UniProt using keywwrds indicating that they do not have antimicrobial activity, and filtered similarly, producing a size- and length-matched set of 4,853 peptides.

We split the AMP and non-AMP sequences using a length-aware, embedding-dissimilar strategy to minimize similarity-driven leakage. The resulting AMP training set contained 8,235 sequences (4,202 positive, 4,033 negative), and the AMP test set contained 1,042 sequences (521 positive, 521 negative). The training set was used to train PLUM, while the held-out test set was used as the prototype set for prototype-guided peptide generation. Full details of the generation of the dataset and its properties are provided in Supplementary Materials Section 1.1.

### PLUM Model Framework

PLUM is a deep generative framework for designing novel AMP sequences. Inspired by conditional subspace VAEs (Klys et al., 2018) and disentangled latent representations in sequential data (Chen et al., 2019; Esmaeili et al., 2019; Mathieu et al., 2019), PLUM encodes peptides into three latent subspaces. The first latent, **Z**_func_, is encouraged to model antimicrobial functionality, allowing explicit control over predicted activity. The second latent, **Z**_length_, is encouraged to capture length-related information, enabling control of peptide size. The third latent, **Z**_res_, is designed to capture residual sequence information not explicitly assigned to AMP activity or length.

By organizing the latent representation around these factors, PLUM supports controllable generation, as each latent component can be modified during sequence design.

The architecture comprises three main components: an LSTM-based encoder that maps peptide sequences to latent representations, structured latent subspaces corresponding to function, length, residual sequence context, and an autoregressive decoder that reconstructs sequences from these latents.

#### Encoding Disentangled Latent Spaces

PLUM maps a peptide sequence **x** into three latent subspaces, **Z**_res_ (residual sequence context), **Z**_func_ (antimicrobial activity), and **Z**_length_ (length bin *ℓ*_bin_).

Each latent is an independent multivariate Gaussian via an LSTM encoder:

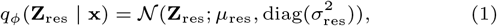

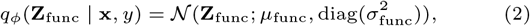

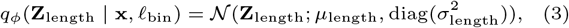

with means *µ*_*k*_ and variances 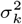 predicted by the encoder (log-variances for stability).

Latents are sampled via the reparameterization trick, **Z**_*k*_ = *µ*_*k*_ + *σ*_*k*_ ⊙ *ϵ*_*k*_, *ϵ*_*k*_ ~ N (0, *I*). This structure supports controlled generation by allowing each latent component to be modified during decoding.

#### Decoder

The PLUM decoder reconstructs peptide sequences from the latent subspaces **Z**_res_, **Z**_func_, and **Z**_length_ using an autoregressive LSTM. At each step, the decoder predicts the next amino acid conditioned on the previously generated residues and the latent codes. For a sequence of length *L*, the generative distribution factorizes as

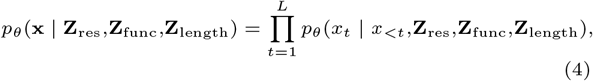

where *x*_*<t*_ = (*x*_1_, …, *x*_*t*−1_). This factorization defines autoregressive sequence generation from the three latent components.

#### Latent Priors

Each latent subspace in PLUM is assigned a prior to structure the latent representation and support conditional control: The sequence latent follows a standard normal,

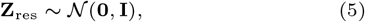

capturing general sequence variability. The functional latent is conditioned on the functional label *y* and follows a Gaussian with parameters predicted by a learned MLP,

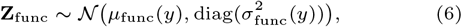

as in the CSVAE framework (Klys et al., 2018). The length latent is conditioned on discrete length bin *b* ∈ {1, …, *B*},

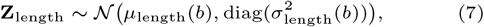

allowing explicit control over peptide length while maintaining variability.

#### Latent Supervision and Adversarial Regularization

PLUM uses auxiliary prediction heads and a secondary decoder to encourage each latent component to capture its intended source of variation. The functional latent predicts antimicrobial activity via

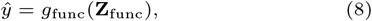

and the length latent predicts the discrete length bin *b*, where

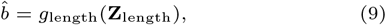

A secondary decoder reconstructs sequences from the sequence latent alone,

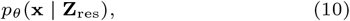

encouraging **Z**_res_ to retain general sequence information. Adversarial predictors attempt to infer *y* and *b* from **Z**_res_, while the encoder is trained to minimize their accuracy. This regularization discourages functional and length information from being encoded in **Z**res, supporting its role as a residual sequence space.

#### Training Objective

PLUM is trained with a multi-objective variational framework combining reconstruction, auxiliary supervision, latent regularization, and adversarial regularization.

##### Sequence reconstruction

The primary decoder (Equation 4) reconstructs the full sequence from all latent subspaces:

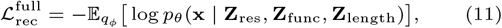

and a secondary decoder reconstructs using only the residual sequence latent (Equation 10):

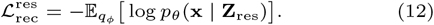

##### Auxiliary supervision

Functional and length latents are regularized via auxiliary prediction heads (Equation 8, Equation 9):

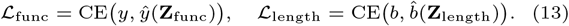

##### Latent regularization

All latent subspaces are regularized via KL divergence from their respective priors (Equation 5 to Equation 7):

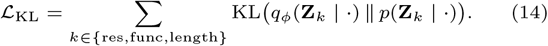

##### Adversarial regularization

Adversarial predictors on the residual sequence latent (Latent Supervision and Adversarial Regularization) attempt to infer *y* and *b*:

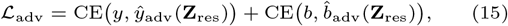

while the encoder is trained to reduce function- and length-predictive information in **Z**_res_.

##### Total loss

The full training objective is

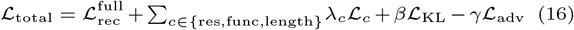

with weights *λ*_res_, *λ*_func_, *λ*_length_, *β, γ* controlling the contribution of each term.

### Training and Generation

PLUM is trained on curated AMP training data using a multi-objective variational framework (subsubsection 2.2.5). The training objective jointly optimizes sequence reconstruction, latent regularization, auxiliary functional and length supervision, and adversarial regularization of the residual sequence latent. Ablation studies showed that the complete training objective and full latent configuration provided the strongest overall balance between reconstruction quality and functional generation performance. Analyses of the individual latent spaces further indicated that class- and length-related information was captured primarily by their corresponding latent components (Supplementary Sections S3.1–S3.3).

For generation, PLUM supports two modes: *de novo* generation, which samples latent variables from activity- and length-conditioned priors to generate sequences autoregressively, and prototype-guided generation, which uses a prototype peptide to bias the decoding process. In prototype-guided generation, the prototype peptide is encoded into the residual sequence latent representation (**Z**_res_), while the functional and length latents (**Z**_func_ and **Z**_length_) are specified to guide the target activity class and peptide length. A tunable prototype-bias parameter (*β*) is applied during token-level decoding, increasing the probability of selecting amino acids consistent with the prototype while allowing alternative sequence compositions. Thus, *β* controls the strength of prototype anchoring, whereas **Z**_func_ and **Z**_length_ guide functional and length-based variation. Details on generation algorithms are provided in Supplementary Material Section S1.4.

### Classifiers for Generated Peptide Prioritization

To prioritize generated AMPs with predicted broad antibacterial activity, we developed the PLUM Antibacterial Potency Model (PLUM-APM). PLUM-APM was trained using MIC-derived labels from an antibacterial potency dataset we developed (More information in supplementary section S1.1.5), containing peptides with activity against both Gram-positive and Gram-negative targets. Peptides with consistent activity across bacterial groups were labeled as high-potency (≤ 10 *µ*M; 927 sequences) or low-potency (*>* 10 *µ*M; 979 sequences), followed by a 75:25 train-test split. The final model used ProtT5 embeddings with a Support Vector Machine (SVM) classifier and was applied to further identify generated peptides predicted as AMPs by external classifiers.

The PLUM software also includes an internal AMP Classifier for optional filtering. The classifier combines ProtT5 embeddings with a Multi-layer Perceptron (MLP) model. Because it was trained on the same curated AMP training set used for PLUM training, it was not used as an external evaluation metric for generated peptides.

The internal AMP classifier achieved performance comparable to the baseline predictor AMPlify, reaching a held-out test accuracy and F1-score of 0.82. For the PLUM-APM model, SVM performed best, achieving 0.85 across all test metrics. Additional dataset construction details and classifier performance results are provided in the Supplementary Materials, Section S1, and Tables S2–S3.

## Results

We evaluated PLUM in *de novo* and prototype-guided generation settings. Generated PLUM sequences were evaluated without post-generation filtering to assess the model’s intrinsic generative capability. AMPlify was used as the primary external AMP classifier, and AMP Scanner V2 was used as a secondary external classifier to verify classification trends (Veltri et al., 2018; Li et al., 2022). We then applied PLUM-APM to estimate antibacterial potency among generated peptides predicted as AMPs.

### Evaluation of De Novo Peptide Generation

PLUM supports function- and length-conditioned *de novo* peptide generation. We evaluated its ability to generate antimicrobial and non-antimicrobial peptides against existing peptide-generation models, focusing on target-class yield, length-wise robustness, and predicted safety and stability.

#### Baselines and evaluators

For benchmarking, PLUM was compared against AMP-GAN (Van Oort et al., 2021), HydrAMP (Szymczak et al., 2023), and two in-house cVAE baselines (Baseline 1 and Baseline 2). These benchmarks were selected due to their ease of use, well-documented implementations, and support for large-scale peptide generation. Baseline 1 uses fully connected encoder–decoder networks, while Baseline 2 uses LSTM-based architectures to capture sequential dependencies. Detailed descriptions of the architectures, training procedures, and generation hyperparameters are provided in the Supplementary Material (Section S2). Functional validity was evaluated using two AMP classifiers, AMP Scanner V2 and AMPlify, with their default parameters. PLUM-APM was then applied in conjunction with AMPlify predictions to identify generated peptides predicted to be both antimicrobial and high-potency against bacterial targets.

#### PLUM achieves a better yield across AMP and non-AMP generation tasks

To generate AMPs and non-AMPs, PLUM was conditioned on a functional latent variable set to 1 (AMPs) or 0 (non-AMPs), with peptide lengths ranging from 5 to 35 amino acids. For HydrAMP, the filter out parameter was disabled to ensure consistent evaluation across models. For each model and functional category, 50,000 peptides were generated, deduplicated, and up to 45,000 unique sequences were retained.

Table 1 summarizes conditioned *de novo* generation performance under AMP and non-AMP settings. PLUM achieved the highest classifier-confirmed yield for both tasks according to AMPlify and AMP Scanner V2. PLUM generated the largest number of peptides predicted to be both antimicrobial by AMPlify and high-potency against bacterial targets by PLUM-APM, indicating a stronger yield of potentially effective antibacterial candidates than the comparison models. HydrAMP, another cVAE-based peptide generator, showed AMP-generation yields comparable to the in-house cVAE baselines. Overall, these results show that PLUM provides effective functional conditioning for both antimicrobial and non-antimicrobial peptide generation.

**Table 1.** Evaluation of *de novo* peptide generation. A total of 45,000 peptides were generated for both AMP and non-AMP categories. Values are reported as percentage yield relative to the 45,000 generated sequences, with the corresponding peptide count in parentheses. AMP and non-AMP yields were evaluated using AMPlify and AMP ScannerV2. Potent AMP yield represents peptides predicted as AMP by AMPlify and as high-potency by the PLUM antibacterial potency model. Toxicity was evaluated using ToxinPred3.

| Generative model | AMP generation task |  |  |  |  | non-AMP generation task |  |  |
| --- | --- | --- | --- | --- | --- | --- | --- | --- |
|  | Total Generated | AMP yield AMPLify | AMP yield ScannerV2 | Potent AMP yield AMPLify+PLUM-APM | Non-toxic yield ToxinPred3 | Generated | non-AMP yield AMPLify | non-AMP yield ScannerV2 |
| AMP-GAN | 45,000 | 49.5 (22,275) | 51.3 (23,063) | 29.2 (13,134) | 74.5 (33,503) | – | – | – |
| HydrAMP | 45,000 | 65.2 (29,318) | 67.2 (30,218) | 45.5 (20,463) | 57.7 (25,971) | 45,000 | 69.6 (31,313) | 72.1 (32,444) |
| Baseline 1 | 45,000 | 68.4 (30,767) | 68.8 (30,975) | 42.6 (19,182) | 74.7 (33,628) | 45,000 | 84.3 (37,935) | 79.2 (35,642) |
| Baseline 2 | 45,000 | 67.8 (30,497) | 68.8 (30,941) | 40.1 (18,047) | **79.9 (35,950)** | 45,000 | 84.8 (38,180) | 78.7 (35,459) |
| **PLUM** | 45,000 | **82.9 (37,324)** | **83.9 (37,754)** | **54.1 (24,358)** | 77.0 (34,650) | 45,000 | **91.9 (41,385)** | **83.6 (37,617)** |

#### PLUM maintains robust generation performance across peptide lengths

We evaluated length-wise generation performance using AMPlify by selecting 300 unique peptides per sequence length, from 5 to 35 amino acids, after deduplication. Diff-AMP (Wang et al., 2024) was included only in the length-wise comparison because its available Hugging Face interface limits generation volume, while direct bulk generation from the released code could not be completed due to an unresolved dependency. Figure 2 shows AMPlify-based target-class yield across peptide lengths for both AMP and non-AMP generation tasks. PLUM showed reduced AMP-generation yield for very short peptides, consistent with the lower representation of short sequences in the training dataset (Supplementary Figure S1). However, performance improved from approximately length 11 onward, remained stable across longer peptides and generated a larger number of predicted AMPs than other models at lengths above 14 AA. In comparison, AMP-GAN showed higher AMP yield at shorter lengths, but this model’s ability to generate AMPs dropped off at lengths above 18 AA. HydrAMP performed similarly to PLUM at shorter lengths and had higher yields at longer lengths, but not to the level of PLUM. Diff-AMP performed the worst out of all models, with the fewest predicted AMPs across all lengths (Figure 2A). For non-AMP generation, PLUM performance exceeded other models across all lengths (Figure 2B).

**Fig. 1.**
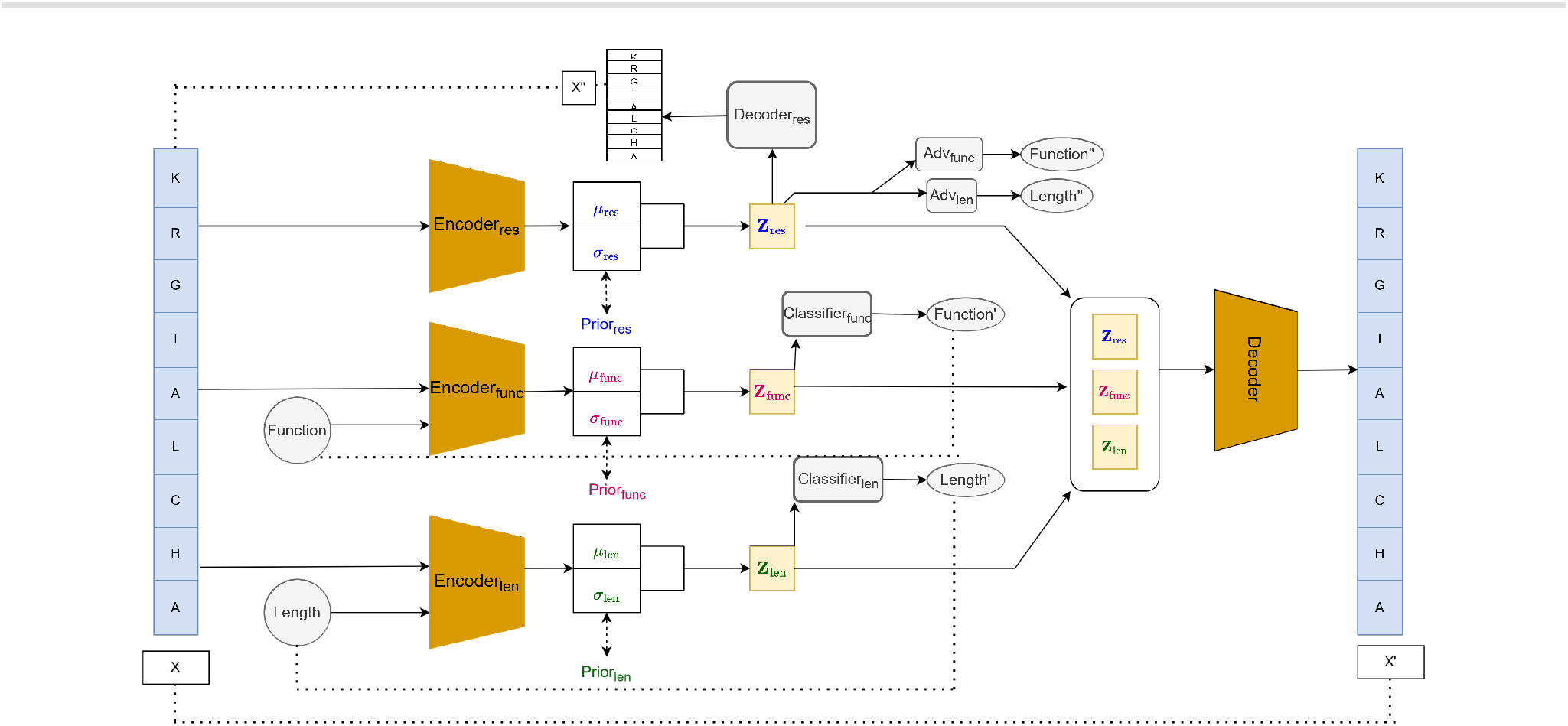
PLUM model architecture. Input peptide *X* is encoded into three latent spaces: *Z*_res_ (residual sequence context), *Z*_func_ (functional activity), and *Z*_length_ (peptide length), each with *µ* and *σ* outputs. KL divergence is computed with priors Prior_res_, Prior_func_, Prior_length_ (*L*_KL_). The main decoder reconstructs *X*^′^ from all latents 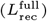, while a secondary decoder reconstructs *X*^′′^ from *Z*_res_ alone 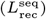. Auxiliary classifiers predict function and length (*L*_func_, *L*_length_), and adversarial heads predict function and length from *Z*_res_ (*L*_adv_). Dotted lines indicate loss computations.

**Fig. 2.**
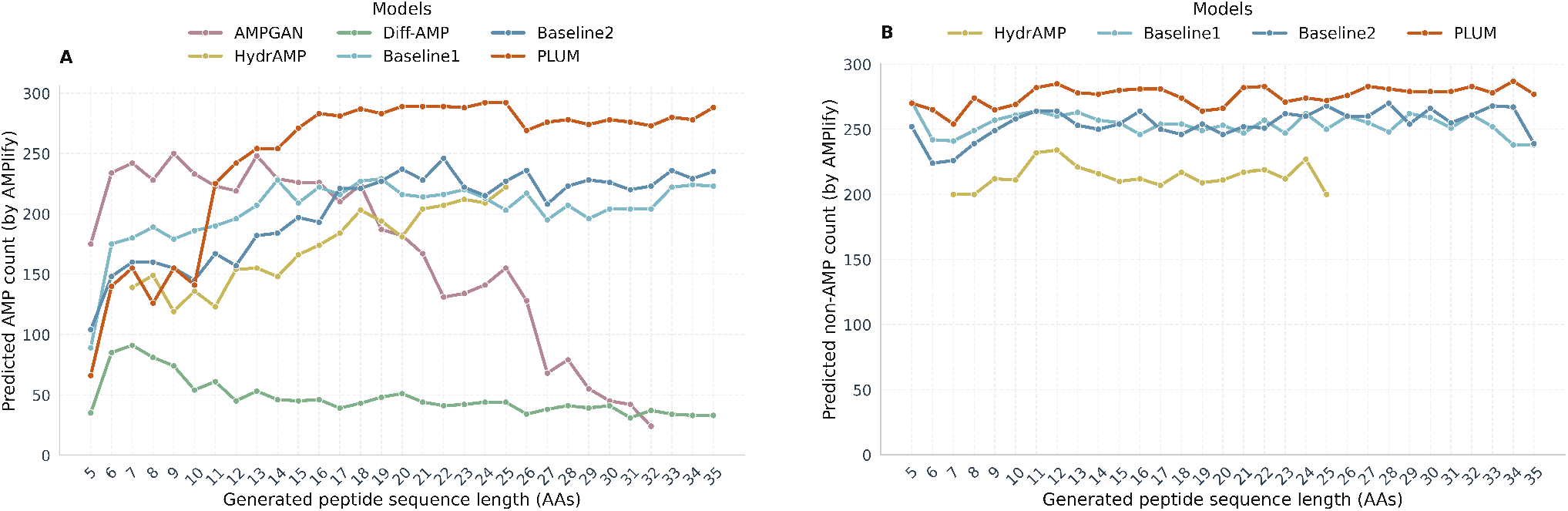
**(A)** AMP generation task. **(B)** non-AMP generation task. Length-wise evaluation of peptide generation performance. For each model and peptide length (5–35 amino acids), 300 sequences were generated and evaluated using AMPlify. The reported counts correspond to peptides predicted as AMPs in the AMP-generation task and peptides predicted as non-AMPs in the non-AMP-generation task.

#### PLUM generates novel and diverse AMPs

We next evaluated whether PLUM-generated AMPs were distinct from the training data and whether the generated set maintained substantial internal diversity. For novelty, each generated AMP was compared with its MMseqs2-selected nearest training-set sequence using normalized Levenshtein similarity and BLOSUM80-based amino-acid similarity. The median nearest-neighbor similarity was 42.86% by normalized Levenshtein similarity and 55.56% by BLOSUM80-based amino-acid similarity, indicating that most generated AMPs remained substantially different from their closest training-set sequences (Supplementary Figure S9).

Internal diversity was assessed by comparing each generated AMP with its nearest non-self generated neighbor using the same two similarity measures. The median nearest-neighbor similarity was 54.5% by normalized Levenshtein similarity and 66.7% by BLOSUM80-based amino-acid similarity, indicating substantial sequence-level variation within the generated set (Supplementary Figure S10). ProtT5-based visualization further showed that PLUM-generated AMPs occupied a broad region of peptide representation space relative to the training and held-out test sets (Supplementary Figure S11). Together, these analyses support that PLUM generates AMPs that are both novel relative to the training data and diverse within the generated population.

#### PLUM generates peptides with favorable predicted safety and stability profiles

We next assessed predicted safety- and stability-associated properties of the generated peptides. PLUM-generated sequences showed favorable predicted toxicity profiles comparable to Baseline 1 and Baseline 2, which may reflect the use of the same training data enriched for lower-toxicity AMP sequences. (Table 1). Their physicochemical property distributions were broadly consistent with known, experimentally validated AMPs (Known AMPs; Section S3.4, Figures S5). Notably, PLUM-generated peptides had the lowest median instability index among the evaluated models (16.40), suggesting favorable predicted sequence stability. Predicted proteolytic resistance of AMPs generated from PLUM across 24 proteases was also comparable to Known AMPs, further supporting the favorable predicted peptide properties (Section S3.4, Figure S7). We also observed that PLUM-generated non-AMPs retained physicochemical properties similar to those of known non-AMPs (Figure S6). As part of this evaluation of PLUM versus other models, we additionally compared AA composition of generated AMPs from PLUM, AMPGAN, HydrAMP and baseline models. Unsurprisingly, we observed that generated AMPs from all models were enriched for Ala, Lys, and Leu, AAs frequently present in AMPs (Decker et al., 2022) and enriched in the reference AMPs dataset (Figure S4). AMPs generated from HydrAMP had higher percentages of Trp and Arg, and lower percentages of Ile and Gly, distinct from other models and the reference dataset (Figure S4). In contrast, PLUM was slightly enriched for Gly relative to other models. These data are likely reflective of the training dataset used to create each model, but HydrAMP in particular stood out as having a bias unique from other models in AA composition in generated AMPs. Together, these analyses suggest that PLUM generates peptides with balanced functional, toxicity, and stability-associated properties.

### Evaluation of Prototype-Guided Peptide Generation

PLUM is designed to generate new peptides from prototype sequences while controlling functional activity through prototype-biased decoding.

#### Experimental Setup

To evaluate the prototype-guided peptide generation, we used the AMP test set as prototypes, comprising 521 sequences labeled as AMP or non-AMP. For PLUM, the prototype sequence was used as a decoding bias, while the functional condition was independently set to generate either AMP or non-AMP peptides. Sequence lengths were sampled within the same length bin of the prototype length, and 100 peptides were generated per prototype under each condition. Prototype anchoring was evaluated using three bias strengths, *β* = 0.45, 0.65, and 0.85, where *β* ranges from no prototype bias (*β* = 0) to strongest prototype bias (*β* = 1). Lower *β* values allow greater sequence exploration, whereas higher *β* values impose stronger bias toward the prototype sequence. Prototype similarity was measured by globally aligning each generated peptide, *y*, to its corresponding starting prototype, *x*, using the BLOSUM80 substitution matrix. The alignment score was normalized by the self-alignment scores of the two sequences:

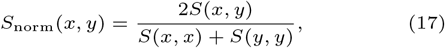

where *S*(*x, y*) is the global BLOSUM80 alignment score between the prototype and generated peptide, and *S*(*x, x*) and *S*(*y, y*) are their respective self-alignment scores. Mean normalized similarity was then calculated across all generated peptides and their corresponding prototypes.

We benchmarked PLUM’s prototype-guided peptide generation approach against two established peptide variant generation methods, HydrAMP and Joker (Porto et al., 2018). For HydrAMP, we used the analogue generation module with n attempts=100 and filtering criteria=‘discovery’. HydrAMP and Joker produced smaller, variable output sets because of their method-specific filtering and constrained analogue-generation procedures. Other baselines were excluded because they do not support prototype-guided generation. Functional validity was assessed using AMPlify, and AMPlify-predicted AMPs were independently screened for predicted antibacterial potency and toxicity using the antibacterial potency classifier and ToxinPred3, respectively.

#### PLUM improves prototype-guided AMP generation over baselines

PLUM generated a substantially larger pool of AMPlify-predicted AMPs than HydrAMP and Joker across both prototype settings (Figure 3). From AMP prototypes, PLUM produced 38,052, 36,465, and 33,730 predicted AMPs at *β* = 0.45, 0.65, and 0.85, respectively, compared with 12,658 for HydrAMP and 3,417 for Joker. PLUM also outperformed both baselines in the more challenging cross-functional setting, generating 23,586, 14,665, and 7,958 predicted AMPs from non-AMP prototypes across the same *β* values, compared with 2,474 for HydrAMP and 1,161 for Joker. PLUM-generated AMP pools retained a substantial subset of AMPs classified as potent by the antibacterial potency model.

**Fig. 3.**
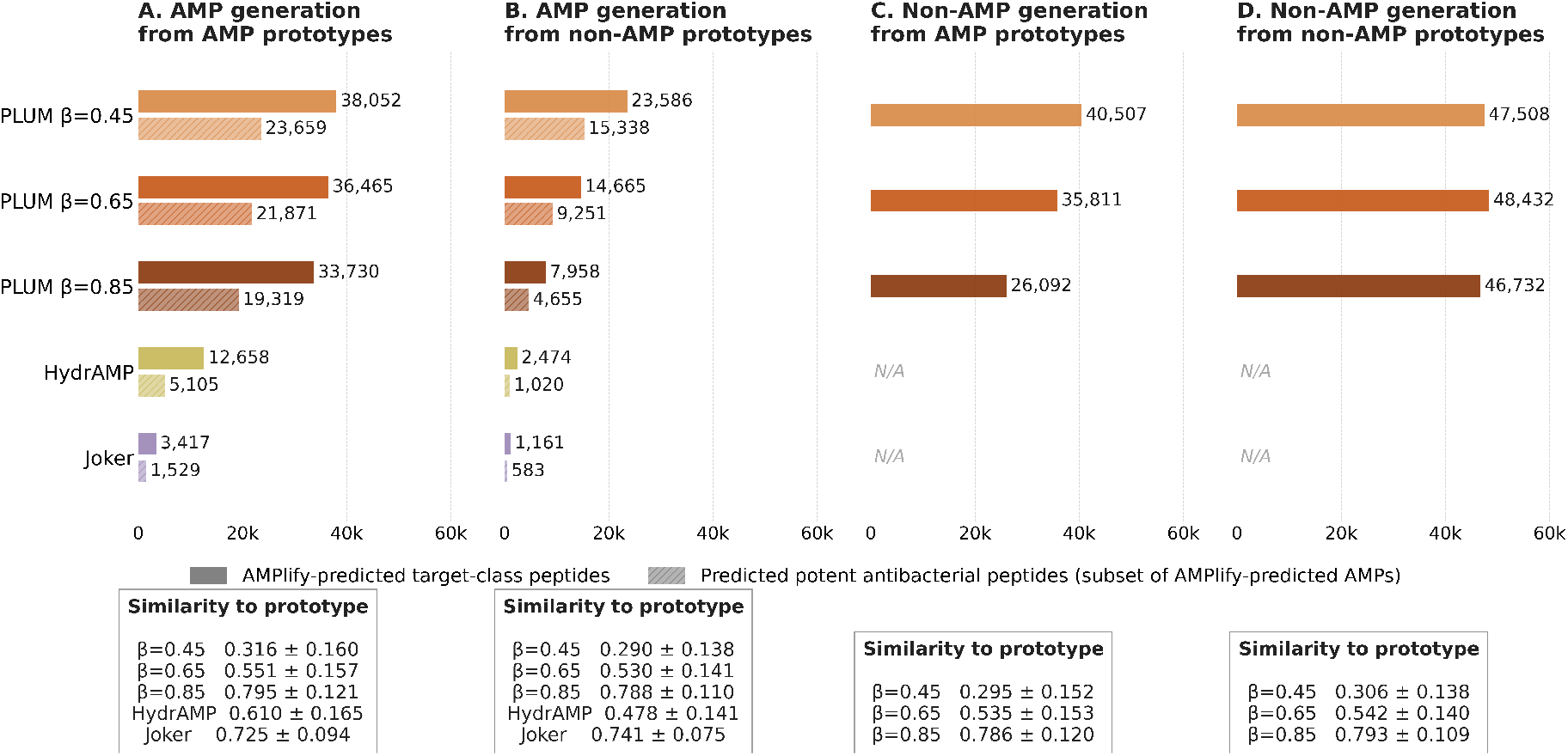
Prototype-guided functional control across AMP and non-AMP generation tasks. The figure compares the number of generated peptides classified as the intended target class by AMPlify. Panels (A) and (B) evaluate AMP generation from AMP and non-AMP prototypes, respectively, while panels (C) and (D) evaluate non-AMP generation from AMP and non-AMP prototypes. For AMP-generation tasks, AMPlify-predicted AMPs were further screened to identify predicted potent antibacterial peptides using the antibacterial potency model (PLUM-APM). PLUM was evaluated using three prototype anchoring strengths (*β* = 0.45, 0.65, and 0.85) and compared against HydrAMP and Joker. HydrAMP and Joker do not support non-AMP-targeted generation; corresponding entries are indicated as unavailable (N/A). Similarity-to-prototype text boxes report the mean *±* SD of the normalized global BLOSUM80 alignment scores between generated peptides and their corresponding starting prototypes.

#### PLUM enables controllable functional exploration across target classes

Beyond AMP generation, PLUM also generated AMPlify-predicted non-AMPs under the non-AMP condition, demonstrating bidirectional functional control. From AMP prototypes, PLUM produced 40,507, 35,811, and 26,092 predicted non-AMPs at *β* = 0.45, 0.65, and 0.85, respectively, while non-AMP prototypes yielded consistently high counts across *β* values: 47,508, 48,432, and 46,732.

The effect of *β* further reflected the tradeoff between functional exploration and prototype anchoring. Consistent with its role as a prototype-anchoring parameter, increasing *β* increased the mean sequence similarity between generated peptides and their paired prototypes across all generation settings. Lower *β* values generally increased target-class yield, whereas higher *β* values imposed stronger prototype bias and reduced functional redirection. This was most evident when generating AMPs from non-AMP prototypes, where AMPlify-predicted AMP counts decreased from 23,586 at *β* = 0.45 to 7,958 at *β* = 0.85. In contrast, when non-AMP prototypes were used to generate non-AMPs, target-class counts remained high across all *β* values, indicating that stronger prototype anchoring was less restrictive when the prototype and target class were aligned.

Additional physicochemical analysis of generated peptides across the four prototype–target settings showed broadly consistent property profiles and is provided in Supplementary Figure S8.

## Discussion

PLUM was developed as a controllable generative framework for designing new antimicrobial peptides and generating prototype-guided peptide variants for therapeutic discovery. A central strength of PLUM is its ability to support large-scale generation of peptides enriched for the intended functional class while retaining control over sequence length. This enables targeted generation of both AMP candidates and non-AMP control sequences across the 5–35 amino acid range. PLUM also showed promising capability of generating lower-toxicity peptides for therapeutic usage, a common issue with AMPs. In *de novo* generation, PLUM improved AMPlify-confirmed AMP yield by 27.3% and non-AMP yield by 32.2% relative to HydrAMP. PLUM also maintained high target-class yield across most peptide lengths from 5 to 35 amino acids, supporting controlled exploration across a range of peptide lengths. Importantly, the generated AMPs also showed low sequence similarity to their nearest training-set neighbors and substantial internal sequence diversity. This indicates that the high functional yield was not limited to generation of sequences closely resembling the training data or to a narrow set of highly similar outputs. Together, this bidirectional functional control and length-wise control enable PLUM to generate AMP candidates and length-matched non-AMP controls for downstream experimental testing and comparison.

Prototype-guided generation extends PLUM beyond *de novo* design by enabling controlled exploration around existing peptide sequences. Using an existing peptide as a prototype, PLUM can generate variants guided by the original sequence while directing generation toward either AMP or non-AMP function. This capability may be useful for expanding peptides with limited target-specific MIC data, exploring improved variants from peptides with moderate activity, and generating matched non-AMP controls for experimental testing.

We believe the addition of the Antibacterial Potency Model provides a practical downstream screening tool for prioritizing AMP candidates. This step is particularly valuable when large candidate sets must be refined for antibacterial applications, including infection control in livestock and clinical healthcare settings.

Overall, PLUM provides a framework for targeted AMP design and establishes a foundation for future model development aimed at improving controllability, flexibility, and candidate prioritization in antimicrobial peptide discovery.

## Data and Code availability

The datasets used during the study are available at: https://doi.org/10.6084/m9.figshare.33158156

The source code used in this study is available at: https://github.com/priyamayur/PLUM

