## Supplemental Materials for "A PLUM Job: Peptide modeLs for Understanding and engineering antiMicrobial therapeutics"

#### S1 PLUM data and model construction

##### S1.1 Extended Dataset Construction

To develop PLUM, we curated standardized datasets of AMPs, non-AMPs, and associated potency measurements. Sequences were collected from public databases and filtered for length, linearity, canonical amino acid composition, sequence validity, and redundancy. The resulting datasets provide a consistent foundation for PLUM training and downstream peptide screening models.

###### S1.1.1 Data sources

AMPs were compiled from four publicly available databases: $CAMP_{R4}$ (released January 2023), DBAASP v3 (released January 2021), DRAMP v4.0 (released January 2025), and GRAMPA (released July 2019)[3, 4, 7, 11]. To construct a corresponding negative dataset of non-antimicrobial peptides, sequences with amino acid lengths between 5 and 35 residues were retrieved from UniProt. Sequences containing any of the following keywords were excluded: "Antimicrobial", "Antibiotic", "Antiviral", "Antifungal", "Fungicide", "Secreted", "Secretory", "Excreted", "Effector", "Defensin", "Disulfide bond", "Cross-link", "Antibacterial", "Bacteriostatic", and "Bactericidal", following the filtering strategy established in [10].

###### S1.1.2 AMP dataset

PLUM is designed to focus on linear AMPs, defined as peptides lacking cyclic or complex structures, disulfide bonds, or non-natural amino acid modifications, and restricted to a length of 5 to 35 amino acids (AA). This design choice reflects the goal of enabling rapid and scalable synthesis of novel PLUM-generated AMPs via solid phase peptide synthesis (SPPS), avoiding inaccessible AMP modifications from a synthesis standpoint [1, 6], and increasing proteolytic resistance and oral bioavailability via use of shorter sequences [9].

Consistent with these design constraints, the positive AMP dataset was compiled by combining sequences from four publicly available AMP databases: CAMP, DRAMP, DBAASP, and GRAMPA. Only natural peptides with lengths between 5 and 35 AA were retained. To exclude cyclic peptides, sequences explicitly labeled as cyclic in DBAASP and DRAMP were removed. Peptides from CAMP and GRAMPA were screened using AlphaFold2 structural predictions to identify and exclude sequences containing cyclic conformations or disulfide bridges. Disulfide bonds were identified by measuring the distance between sulfur atoms of cysteine residues; cysteine pairs with S–S distances below 2.5 Å were considered disulfide-linked. Cyclic peptides were detected by measuring the distance between the N- and C-terminal alpha carbons; sequences with head-to-tail distances below 2 Å were treated as cyclic [2, 12]. Following these filtering steps, the datasets contained 3,625, 2,888, 1,008, and 1,681 sequences from CAMP, DRAMP, DBAASP, and GRAMPA, respectively.

The combined set of 9,202 sequences was subsequently screened against the UniProt database to identify peptides with annotated post-translational modifications (PTMs), and sequences with positive PTM records were removed. The resulting dataset was then further filtered to exclude peptides containing non-canonical amino acids and duplicate entries, yielding a final dataset of 4,723 unique linear AMP sequences.

###### S1.1.3 Non-AMP dataset

The negative dataset was filtered similarly, retaining sequences with lengths between 5 and 35 AAs and only including entries with protein- or transcript-level evidence in UniProt to ensure reliability. Linear peptides were identified using AlphaFold2, applying the same criteria for linearity as described for the AMP dataset, yielding 9,917 sequences. Sequences with annotated post-translational modifications (PTMs) were then removed based on UniProt records. Further filtering for valid amino acids and uniqueness produced 9,090 sequences. To create a balanced dataset, the negative set was adjusted to match both the size and length distribution of the positive set, resulting in a final set of 4,853 sequences (see Supplementary Material Figure S1).

###### S1.1.4 AMP Training and Test Dataset

We combined the curated AMP and non-AMP datasets to construct training and held-out test sets for PLUM. To reduce similarity-driven data leakage, sequences were split using a length-aware, embedding-dissimilar strategy. Peptides were first grouped by sequence length to preserve length distributions. Within each length group, candidate test sequences were selected and compared against candidate training sequences using ProtT5 embeddings and cosine similarity. Test candidates were retained only if their similarity to all training candidates fell below a threshold defined by the $98.4^{\text{th}}$ percentile of pairwise AMP sequence similarities. This threshold was selected to reduce train–test similarity while retaining an approximately balanced test set of 500 sequences per class.

The final split produced 8,235 training sequences (4,202 AMP and 4,033 non-AMP) and 1,042 held-out test sequences (521 AMP and 521 non-AMP). The difference between the total number of curated non-AMPs and the final split reflects the exclusion of some non-AMP sequences required to maintain length-aware, embedding-dissimilar, and class-balanced partitions.

###### S1.1.5 Antibacterial MIC Dataset

To construct the dataset for the Antibacterial Potency Classifier, we compiled experimentally reported minimum inhibitory concentration (MIC) values for AMPs from CAMP, DRAMP, DBAASP, and GRAMPA. MIC values were standardized to micromolar units ($\mu$M), and peptides containing non-canonical amino acids were removed.

For each unique peptide–organism pair, multiple reported MIC measurements were collapsed into a single target-wise consensus MIC value. When two MIC measurements were available, the pair was retained only if the values differed by no more than 5 $\mu$M, and the consensus MIC was calculated as their arithmetic mean. When three or more measurements were available, the consensus MIC was calculated using the geometric mean.

To define sequence-level antibacterial potency labels, we retained only peptides with MIC measurements against at least one Gram-positive and one Gram-negative bacterial species. For each retained peptide, the difference between the highest and lowest target-wise consensus MIC values was required to be no greater than 5 $\mu$M, removing peptides with large variation in activity

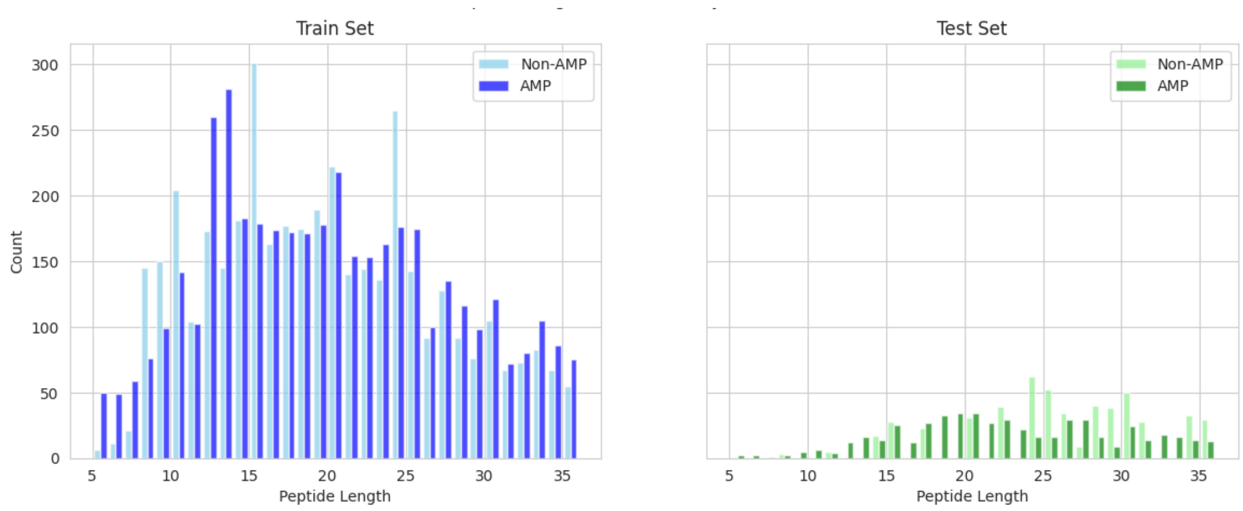


Figure S1: Distribution of AMPs and non-AMPs across different lengths in Train and Test datasets.

across bacterial targets. The final MIC assigned to each peptide was calculated as the mean of its target-wise consensus MIC values.

Peptides with final MIC values $\leq 10$ $\mu$M were labeled as high-potency antibacterial peptides, whereas those with MIC values $> 10$ $\mu$M were labeled as low-potency antibacterial peptides, based on experimental thresholds reported in the literature [13, 5, 8]. The final dataset contained 927 high-potency and 979 low-potency peptides and was split into training and test sets using a 75/25 split.

#### S1.2 Training Hyperparameters

PLUM is trained using a multi-objective conditional VAE framework with LSTM-based encoders and decoders. The training hyperparameters are summarized as follows:

- **Optimizer:** Adam
- **Learning rate:** 0.001
- **Batch size:** 64
- **Number of epochs:** 150 max (early stopping)
- **Hidden dimension:** 128
- **Latent dimensions:** $z = 8$, $w = 8$, $v = 8$
- **Conditioning dimension:** 1
- **Maximum sequence length:** 35

**Loss weights.** The multi-objective loss combines several terms:

- length_loss_weight = 2.0 (length reconstruction)
- func_loss_weight = 2.0 (functional classification)
- z_rec_weight = 1.0 (z-only reconstruction)
- kl_z_weight = 1 (KL divergence for residual sequence latent)
- kl_w_weight = 0.03 (KL divergence for function latent)
- kl_v_weight = 0.03 (KL divergence for length latent)
- adv_weight = 1.0 (adversarial disentanglement)

**Training details.** The decoder uses teacher forcing during training (`teacher_forcing=True`). KL divergence is applied to all latent variables ($z$, $w$, $v$) relative to their learned priors. The multi-objective loss also includes sequence reconstruction, z-only reconstruction, functional classification, length reconstruction, and adversarial disentanglement terms.

#### S1.3 Implementation Details

##### Software and Frameworks

PLUM was implemented in Python 3.10 using PyTorch 2.1. The LSTM encoder and decoder architectures were built with PyTorch's native modules. Auxiliary and adversarial heads were implemented as simple feedforward networks (MLPs) with ReLU activations. Training scripts, data preprocessing, and generation pipelines leverage NumPy, pandas, and standard scientific computing libraries.

##### Hardware

All experiments were performed on NVIDIA A100 GPUs with 40 GB memory. Model checkpointing and logging were handled using standard PyTorch utilities.

##### Data Preprocessing

Peptide sequences were one-hot encoded including `START`, `STOP`, and `PAD` tokens. Sequence lengths were discretized into $B$ predefined bins. Functional labels (AMP vs non-AMP) were encoded as integer classes. Sequences exceeding the maximum length or containing non-standard amino acids were filtered out.

##### Model Architecture Details

The LSTM encoder and decoder each consist of 2 layers with 128 hidden units. Latent subspaces ($\mathbf{Z}_{\text{res}}$, $\mathbf{Z}_{\text{func}}$, $\mathbf{Z}_{\text{length}}$) are 8-dimensional each. Auxiliary and adversarial heads are 2-layer MLPs with 128 hidden units.

##### Training Procedure

Models were trained using the Adam optimizer with a learning rate of $1\times10^{-3}$ and a batch size of 64. Training ran for 150 epochs with early stopping, depending on the experiment. The decoder uses teacher forcing (`teacher_forcing=True`). KL divergence is applied to all latent variables ($z$, $w$, $v$) relative to their learned priors. The multi-objective loss combines sequence reconstruction, z-only reconstruction, functional classification, length reconstruction, and adversarial disentanglement terms. Hyperparameters for the loss weights were selected empirically.

##### Generation Pipeline

For generation, latent variables are sampled from their respective priors. Prototype-conditioned generation perturbs the sequence latent around an existing peptide, while functional and length latents are set to desired values. The decoder generates sequences autoregressively, and sequences are filtered to ensure valid amino acid composition and adherence to target length bins.

#### S1.4 Generation Algorithms

PLUM supports two primary modes for generating peptide sequences: **De novo generation** and **Prototype-conditioned generation**. Both modes leverage the structured latent spaces to produce sequences with controllable functional and structural properties.

##### De Novo Generation

In de novo generation, sequences are produced without requiring a starting peptide. Latent variables are sampled from their respective priors: the residual sequence latent $\mathbf{Z}_{\text{res}}$, the functional latent $\mathbf{Z}_{\text{func}}$, and the length latent $\mathbf{Z}_{\text{length}}$. Optional constraints on functional activity or length bins can be applied by conditioning the corresponding latent variables. Sequences are then generated autoregressively using the decoder. This mode enables exploration of novel peptide sequence space without relying on any existing peptide template.

**Algorithm 1** De novo Peptide Generation

**Require:** Latent priors $p(\mathbf{Z}_{\text{res}})$, $p(\mathbf{Z}_{\text{func}})$, $p(\mathbf{Z}_{\text{length}})$, functional label $y$ and length bin $b$ for length $l$, decoder $p_\theta$

**Ensure:** Generated peptide sequence $\mathbf{x}$

Sample sequence latent: $\mathbf{Z}_{\text{res}} \sim p(\mathbf{Z}_{\text{res}})$
Sample functional latent: $\mathbf{Z}_{\text{func}} \sim p(\mathbf{Z}_{\text{func}} \mid y)$
Sample length latent: $\mathbf{Z}_{\text{length}} \sim p(\mathbf{Z}_{\text{length}} \mid b)$
Initialize empty sequence: $\mathbf{x} \leftarrow$ START token
**while** not $l$ reached **do**
    Compute decoder logits: $\mathbf{l}_t = p_\theta(\mathbf{x}_{<t}, \mathbf{Z}_{\text{res}}, \mathbf{Z}_{\text{func}}, \mathbf{Z}_{\text{length}})$
    Optionally scale logits by temperature: $\mathbf{l}_t \leftarrow \mathbf{l}_t / T$
    Sample next amino acid: $x_t \sim \text{Softmax}(\mathbf{l}_t)$
    Append $x_t$ to sequence $\mathbf{x}$
**end while**
**return x**

##### Prototype-Guided Generation

This algorithm generates peptide variants from a prototype sequence. The prototype is encoded into the residual sequence latent $\mathbf{Z}_{\text{res}}$, with optional Gaussian perturbation controlled by $\sigma$ to explore nearby variants. The functional latent $\mathbf{Z}_{\text{func}}$ specifies the target activity, while $\mathbf{Z}_{\text{length}}$ controls the target length. During autoregressive decoding, the prototype-bias parameter $\beta$ shifts amino acid sampling toward the prototype sequence, balancing prototype similarity with sequence diversity. Hard length constraints further control sequence stochasticity.

**Algorithm 2** Prototype-Guided Peptide Generation

**Require:** Prototype peptide $\mathbf{x}_{\text{proto}}$, latent encoder $q_\phi$, functional label $y$,length bin $b$ for length $l$, perturbation magnitude $\sigma$, soft bias $\beta$, min/max lengths, decoder $p_\theta$

**Ensure:** Generated peptide sequence $\mathbf{x}$

Encode prototype: $\mathbf{Z}_{\text{res}}^{\text{proto}} \sim q_\phi(\mathbf{Z}_{\text{res}} \mid \mathbf{x}_{\text{proto}})$
Optionally perturb: $\mathbf{Z}_{\text{res}} \leftarrow \mathbf{Z}_{\text{res}}^{\text{proto}} + \epsilon$, $\epsilon \sim \mathcal{N}(0, \sigma^2)$
Set functional latent: $\mathbf{Z}_{\text{func}} \sim p(\mathbf{Z}_{\text{func}} \mid y)$
Set length latent: $\mathbf{Z}_{\text{length}} \sim p(\mathbf{Z}_{\text{length}} \mid b)$
Initialize sequence with START token: $\mathbf{x} \leftarrow$ START
**while** not $l$ reached **do**
    Compute decoder probabilities: $\mathbf{p}_t = p_\theta(x_t \mid \mathbf{x}_{<t}, \mathbf{Z}_{\text{res}}, \mathbf{Z}_{\text{func}}, \mathbf{Z}_{\text{length}})$
    Apply soft prototype bias: $\mathbf{p}_t \leftarrow \beta \cdot \mathbf{x}_t^{\text{proto}} + (1 - \beta) \cdot \mathbf{p}_t$
    Sample or select next amino acid $x_t$ from $\mathbf{p}_t$
    Append $x_t$ to sequence $\mathbf{x}$
**end while**
**return x**

Key hyperparameters used in our experiments included a perturbation magnitude $\sigma = 0$, prototype bias $\beta$ was varied for our experiments, minimum and maximum lengths $[5, 35]$, and a sampling temperature $T = 1.0$. Stochastic sampling was enabled to allow diverse sequence generation.

### S2 Baseline Methods

#### S2.1 Baseline 1: Conditional Sequence Variational Autoencoder (cVAE)

This baseline employs a conditional variational autoencoder (cVAE) to generate peptide sequences conditioned on functional activity and sequence length. Peptides are represented as one-hot encoded sequences, with variable lengths handled via masking. Conditioning variables are explicitly incorporated into both the encoder and decoder.

##### S2.1.1 Model Architecture

The model consists of a feedforward encoder–decoder architecture. The encoder maps an input peptide sequence $\mathbf{x}$ and a conditioning vector $\mathbf{c}$ to a latent Gaussian distribution parameterized by a mean $\boldsymbol{\mu}$ and log-variance $\boldsymbol{\log \sigma^2}$. The conditioning vector comprises two components: (i) a binary functional label and (ii) a normalized peptide length.

The encoder flattens the one-hot encoded peptide sequence and concatenates it with the conditioning vector before passing it through fully connected layers with ReLU activations. The decoder mirrors this structure, taking a sampled latent vector $\mathbf{z}$ concatenated with the same conditioning vector and mapping it to a sequence of amino acid logits. The output is reshaped to produce position-wise categorical distributions over the amino acid vocabulary, with padding positions masked during training.

##### S2.1.2 Latent Variable Formulation

Latent variables are sampled using the reparameterization trick:

$$\mathbf{z} = \boldsymbol{\mu} + \boldsymbol{\epsilon} \odot \boldsymbol{\sigma}, \quad \boldsymbol{\epsilon} \sim \mathcal{N}(\mathbf{0}, \mathbf{I}).$$

For prototype-conditioned generation, the latent mean $\boldsymbol{\mu}$ of a peptide is used deterministically, and small Gaussian perturbations are applied to explore structurally related variants.

##### S2.1.3 Training Objective

The model is trained by minimizing a weighted sum of a reconstruction loss and a Kullback–Leibler (KL) divergence term:

$$\mathcal{L} = \mathcal{L}_{\text{rec}} + \lambda \, \mathcal{L}_{\text{KL}},$$

where $\lambda = 0.01$ in our experiments. The reconstruction loss is a categorical cross-entropy over amino acids at each sequence position, ignoring padded positions. The KL divergence regularizes the latent space toward a standard normal prior.

##### S2.1.4 De Novo Peptide Generation

Peptides are generated *de novo* by sampling latent vectors from the prior distribution and decoding them under specified functional and length conditions. The decoded sequences are truncated to the desired length, and amino acids are sampled independently at each position from the softmax-normalized outputs. The temperature scale was set to 1 for our experiments.

**Algorithm 3** De Novo Peptide Generation with cVAE

**Require:** Trained decoder $D$, latent dimension $L$, target length $s$, functional label $f$, temperature $T$ set to 1

Sample latent vector $z \sim \mathcal{N}(0, I^L)$

Create conditioning vector $c = [f, s/\text{max\_seq\_len}]$

Decode logits: $x_{\text{logits}} = D(z, c)$

Truncate logits to target length: $x_{\text{logits}} = x_{\text{logits}}[: s, :]$

Apply softmax with temperature $T$ and sample amino acids: $x \sim \text{Softmax}(x_{\text{logits}}/T)$

**return** generated peptide sequence $x$

**Model, Training, and Generation Hyperparameters** The cVAE model was trained with a latent dimension of $L = 8$ and hidden layers of 128 units in both the encoder and decoder. Peptides were represented as one-hot encoded sequences of 20 amino acids, with variable-length sequences padded to the maximum length in the dataset. The model was optimized using the Adam optimizer with a learning rate of $1 \times 10^{-3}$ and a batch size of 16 over 150 epochs. The training objective consisted of a masked categorical cross-entropy reconstruction loss and a KL divergence term weighted by $\lambda = 0.01$.

During peptide generation, conditioning vectors combined the functional label and normalized sequence length. De novo peptides were generated by sampling latent vectors from the prior with temperature $T = 1.0$, whereas prototype-conditioned analogues were generated by encoding the prototype, adding Gaussian perturbations to the latent vector with standard deviation $\sigma = 0.01$, and decoding under the target function and length with temperature $T = 1.0$. Amino acids were sampled independently at each sequence position for both generation modes.

#### S2.2 Baseline 2: Conditional Sequence Variational Autoencoder with LSTM (cVAE LSTM)

This baseline employs a conditional variational autoencoder (cVAE) with recurrent neural networks to generate peptide sequences. Peptides are represented as one-hot encoded sequences augmented with special tokens for start-of-sequence (SOS), end-of-sequence (EOS), and padding (PAD). Variable-length sequences are handled via padding and packed LSTM sequences. Conditioning variables, comprising the functional label and normalized peptide length, are incorporated into both the encoder and decoder.

##### S2.2.1 Model Architecture

The encoder is a single-layer LSTM that processes packed peptide sequences. The final hidden state is concatenated with the conditioning vector and passed through linear layers to produce the latent mean $\boldsymbol{\mu}$ and log-variance $\log \boldsymbol{\sigma}^2$. The decoder is an autoregressive LSTM with an embedding layer, which generates one token at a time. Its initial hidden state is computed from the concatenation of the latent vector and conditioning vector. At each time step, the decoder outputs logits over the amino acid vocabulary plus special tokens, and the next token is sampled either from the target sequence (teacher forcing) or from the decoder output.

##### S2.2.2 Latent Variable Formulation

Latent variables are sampled using the reparameterization trick:

$$\mathbf{z} = \boldsymbol{\mu} + \boldsymbol{\epsilon} \odot \boldsymbol{\sigma}, \quad \boldsymbol{\epsilon} \sim \mathcal{N}(\mathbf{0}, \mathbf{I}).$$

During prototype-conditioned generation, the latent mean $\boldsymbol{\mu}$ is used deterministically, with small Gaussian perturbations applied to generate structurally related analogues.

##### S2.2.3 Training Objective

The model is trained by minimizing a weighted sum of a masked categorical cross-entropy reconstruction loss and a KL divergence regularization term:

$$\mathcal{L} = \mathcal{L}_{\text{rec}} + \lambda\,\mathcal{L}_{\text{KL}},$$

with $\lambda = 0.01$. The reconstruction loss ignores padded positions. Teacher forcing is applied during training, with a ratio decaying from 1.0 to 0.1 over epochs. The model is optimized using the Adam optimizer with a learning rate of $1 \times 10^{-3}$, a batch size of 16, and trained for 500 epochs.

##### S2.2.4 De Novo Peptide Generation

Peptides are generated *de novo* by sampling latent vectors from the prior distribution and decoding them autoregressively under specified functional and length conditions. Generation stops when either the EOS token is sampled or the target length is reached. The temperature scale was set to 1 for all our experiments.

**Algorithm 4** De Novo Peptide Generation with cVAE LSTM

**Require:** Trained decoder $D$, latent vector $z$, target length $s$, functional label $f$, temperature $T$

1: Create conditioning vector $c = [f, s/D.max\_len]$
2: Decode sequence autoregressively with decoder $D(z, c)$
3: Truncate to target length or stop at EOS token
4: Apply softmax with temperature $T$ and sample amino acids
5: **return** generated peptide sequence

**Model, Training, and Generation Hyperparameters** The cVAE LSTM was trained with a latent dimension of $L = 16$, LSTM hidden size of 128, and embedding dimension of 64 in the decoder. Peptides were represented with one-hot encoding over 20 amino acids plus SOS, EOS, and PAD tokens, padded to the maximum sequence length. The optimizer was Adam with a learning rate of $1 \times 10^{-3}$, batch size 16, over 150 epochs. The KL divergence weight was $\lambda = 0.01$, and teacher forcing decayed from 1.0 to 0.1 during training.

During generation, conditioning vectors combined the functional label and normalized sequence length. De novo peptides were generated with temperature $T = 1.0$, while prototype-conditioned analogues used Gaussian perturbations with $\sigma = 0.01$ and temperature $T = 1.0$. Amino acids were sampled independently at each step, stopping at EOS or the target length.

Table S1: Comparison of Baseline 1 and Baseline 2 cVAE models

| **Feature** | **Baseline 1** | **Baseline 2** |
|---|---|---|
| Encoder | Fully connected feedforward | Single-layer LSTM (packed sequences) |
| Decoder | Fully connected feedforward | Autoregressive LSTM with embedding |
| Sequence representation | One-hot (20 AAs) | One-hot + SOS/EOS/PAD (23 tokens) |
| Conditioning | Functional label + normalized length | Functional label + normalized length |
| Latent dimension | 8 | 16 |
| KL weight ($\lambda$) | 0.01 | 0.01 |
| Teacher forcing | N/A | Decaying from 1.0 $\rightarrow$ 0.1 |
| Generation mode | Parallel, per-position | Stepwise autoregressive, stop at EOS or target length |
| Prototype-conditioned generation | Add Gaussian noise ($\sigma = 0.01$) | Add Gaussian noise ($\sigma = 0.01$) |
| Training epochs | 500 | 500 |
| Batch size | 16 | 16 |
| Optimizer | Adam (LR $1 \times 10^{-3}$) | Adam (LR $1 \times 10^{-3}$) |

#### S3 Supplementary Analysis

Table S2: **Performance of the AMP Classifier.** Accuracy, precision, recall, and F1-score are reported for training/CV and held-out test data. For ProtT5-based models, training/CV results represent mean ± standard deviation across cross-validation folds, while held-out test results are reported as weighted averages. External AMP predictors are included for comparison on the same datasets. Best values in each column are shown in **bold**.

| Model | Train / CV (mean ± std) | | | | Test | | | |
|---|---|---|---|---|---|---|---|---|
| | Accuracy | Precision | Recall | F1-score | Accuracy | Precision | Recall | F1-score |
| **ProtT5 + MLP** | 0.884 ± 0.019 | 0.884 ± 0.019 | 0.884 ± 0.019 | 0.884 ± 0.019 | **0.820** | **0.840** | **0.820** | **0.820** |
| ProtT5 + SVM | 0.893 ± 0.006 | 0.897 ± 0.006 | 0.894 ± 0.006 | 0.893 ± 0.006 | 0.810 | **0.840** | 0.810 | 0.810 |
| ProtT5 + RF | 0.884 ± 0.019 | 0.885 ± 0.019 | 0.884 ± 0.019 | 0.884 ± 0.019 | 0.770 | 0.810 | 0.770 | 0.770 |
| AMPLIFY | 0.870 | 0.870 | 0.870 | 0.870 | **0.820** | 0.810 | 0.810 | 0.810 |
| DIFF-AMP | 0.810 | 0.810 | 0.810 | 0.810 | 0.720 | 0.750 | 0.720 | 0.710 |
| Peptide Scanner | 0.830 | 0.830 | 0.830 | 0.830 | 0.790 | 0.800 | 0.790 | 0.790 |

Table S3: **Performance of the Antibacterial Potency Classifier.** Accuracy, precision, recall, and F1-score are reported for cross-validation and held-out test data. Cross-validation values are mean ± standard deviation across folds, while held-out test metrics are reported as weighted averages. Best values in each column are shown in **bold**.

| Model | Cross validation (mean ± std) | | | | Test | | | |
|---|---|---|---|---|---|---|---|---|
| | Accuracy | Precision | Recall | F1-score | Accuracy | Precision | Recall | F1-score |
| **ProtT5 + SVM** | **0.8322 ± 0.0241** | **0.8326 ± 0.0242** | **0.8322 ± 0.0241** | **0.8322 ± 0.0241** | **0.85** | **0.85** | **0.85** | **0.85** |
| ProtT5 + RF | 0.8257 ± 0.0235 | 0.8260 ± 0.0232 | 0.8257 ± 0.0235 | 0.8255 ± 0.0237 | 0.83 | 0.83 | 0.83 | 0.83 |
| ProtT5 + MLP | 0.8289 ± 0.0282 | 0.8298 ± 0.0281 | 0.8289 ± 0.0282 | 0.8289 ± 0.0282 | 0.83 | 0.83 | 0.83 | 0.83 |

Table S4: **Peptide generation efficiency across models.** Wall-clock time required by each generative model to produce 50,000 peptides on resource-constrained, CPU-only hardware with 8 GB RAM.

| Model | Generation settings | Time |
|---|---|---|
| AMPGAN | Default configuration | ∼30 min |
| HydrAMP | Default (`properties=True`, `filter_out=True`, `n_attempts=64`) | ∼3.5 min |
| HydrAMP | User-defined (`properties=False`, `filter_out=False`, `n_attempts=1`) | ∼4 s |
| PLUM | Default configuration | ∼3 s |

#### S3.1 Ablation of Training Loss Components

To evaluate the contribution of individual training losses to PLUM's performance, we conducted an ablation study in which specific components of the full loss function were removed. The full training loss is defined as:

$$\mathcal{L}_{\text{total}} = \mathcal{L}_{\text{rec}}^{\text{full}} + \lambda_{\text{res}}\mathcal{L}_{\text{rec}}^{\text{res}} + \lambda_{\text{func}}\mathcal{L}_{\text{func}} + \lambda_{\text{length}}\mathcal{L}_{\text{length}} + \beta\mathcal{L}_{\text{KL}} - \gamma\mathcal{L}_{\text{adv}}. \tag{1}$$

We systematically removed individual terms to assess their impact on peptide generation quality. Generated sequences were evaluated using PLUM's internal AMP classification model, reporting Accuracy, F1 score, Precision, and Recall. Additionally, reconstruction loss ($\mathcal{L}_{\text{rec}}^{\text{full}}$) was recorded as a measure of sequence fidelity. The results of this ablation study are summarized in Table S5, showing the impact of removing individual loss components on sequence reconstruction and AMP classification performance.

Table S5: Ablation study of PLUM training losses. Metrics were computed using the internal AMP classifier on 10,000 generated sequences per functional condition.

| **Loss Configuration** | **Reconstruction loss** | **Accuracy** | **F1** | **Precision** | **Recall** |
|---|---|---|---|---|---|
| Full model | 1.5594 | 0.9086 | 0.9076 | 0.9178 | 0.8976 |
| No z-only decoder | 1.8263 | 0.8569 | 0.8557 | 0.8634 | 0.8480 |
| No adversarial | 1.6541 | 0.9071 | 0.9042 | 0.9335 | 0.8767 |
| No z-only decoder + No adversarial | 1.6826 | 0.8691 | 0.8623 | 0.9099 | 0.8194 |
| No functional loss | 1.5567 | 0.8547 | 0.8509 | 0.8739 | 0.8291 |
| No length loss | 1.6180 | 0.8256 | 0.8092 | 0.8928 | 0.7400 |
| No functional & length loss | 1.5543 | 0.8467 | 0.8403 | 0.8768 | 0.8067 |

#### S3.2 Ablation of Latent Subspaces

To assess the contribution of each latent subspace in PLUM, we performed an ablation study comparing models trained with selective combinations of latent variables:

- **Z + W only:** Retains sequence ($\mathbf{Z}_{\text{res}}$) and functional ($\mathbf{Z}_{\text{func}}$) latents; length latent excluded.
- **Z + V only:** Retains sequence ($\mathbf{Z}_{\text{res}}$) and length ($\mathbf{Z}_{\text{length}}$) latents; functional latent excluded.
- **Full Z + W + V:** Baseline model with all three latent subspaces.

Generated sequences from each configuration were evaluated using reconstruction loss ($\mathcal{L}_{\text{rec}}^{\text{full}}$) and classification metrics (Accuracy, F1, Precision, Recall), consistent with the evaluation in the training loss ablation study. The results are summarized in Table S6.

Table S6: Ablation of latent subspaces. Excluding specific latents affects functional or length control.

| Latent Configuration | Reconstruction loss | Accuracy | F1 | Precision | Recall |
|---|---|---|---|---|---|
| Z + W only | 1.5234 | 0.8471 | 0.8415 | 0.8732 | 0.8120 |
| Z + V only | 1.5564 | 0.8689 | 0.8665 | 0.8825 | 0.8510 |
| Full Z + W + V | 1.5594 | 0.9086 | 0.9076 | 0.9178 | 0.8976 |

Excluding the functional latent ($\mathbf{Z}_{\text{func}}$) reduces functional specificity, while removing the length latent ($\mathbf{Z}_{\text{length}}$) leads to poorer length control. The residual sequence latent ($\mathbf{Z}_{\text{res}}$) is essential for high-fidelity sequence reconstruction. Overall, the full Z + W + V configuration achieves the best balance of reconstruction accuracy and functional predictability, supporting that all three latent subspaces provide complementary information critical for interpretable and controllable peptide generation.

Table S7: **Evaluation of information captured by individual PLUM latent subspaces.** Prediction models were trained using each latent subspace separately and evaluated on the held-out test set. Binary class-label prediction and continuous length prediction were used to assess how much class- and length-related information each subspace captured.

| **Binary class label prediction** | | | | | |
|---|---|---|---|---|---|
| Latent component | Accuracy | Precision | Recall | Specificity | F1 |
| $\mathbf{Z}_{\text{res}}$ | 44.82 | 44.94 | 46.07 | 43.57 | 45.50 |
| $\mathbf{Z}_{\text{func}}$ | 100.00 | 100.00 | 100.00 | 100.00 | 100.00 |
| $\mathbf{Z}_{\text{length}}$ | 54.03 | 52.43 | 86.95 | 21.11 | 65.42 |
| $\mathbf{Z}_{\text{res}} + \mathbf{Z}_{\text{func}} + \mathbf{Z}_{\text{length}}$ | 100.00 | 100.00 | 100.00 | 100.00 | 100.00 |

| **Continuous length prediction** | | | |
|---|---|---|---|
| Latent component | MAE | MSE | $R^2$ |
| $\mathbf{Z}_{\text{res}}$ | 6.585 | 62.659 | -47.33 |
| $\mathbf{Z}_{\text{func}}$ | 6.753 | 65.159 | -53.21 |
| $\mathbf{Z}_{\text{length}}$ | 0.270 | 0.180 | 99.58 |
| $\mathbf{Z}_{\text{res}} + \mathbf{Z}_{\text{func}} + \mathbf{Z}_{\text{length}}$ | 0.508 | 0.341 | 99.20 |

#### S3.3 Evaluating individual latent subspaces

To evaluate the role of each latent subspace, we trained MLP-based predictive models using each latent component separately and evaluated them on the held-out test set (Table S7). The function-associated component, $\mathbf{Z}_{\text{func}}$, accurately predicted the binary class label, while $\mathbf{Z}_{\text{res}}$ and $\mathbf{Z}_{\text{length}}$ performed much worse for this task. In contrast, the length-associated component, $\mathbf{Z}_{\text{length}}$, was the best predictor of peptide length, with a mean absolute error of 0.270 amino acids and an $R^2$ of 99.58%. $\mathbf{Z}_{\text{res}}$ and $\mathbf{Z}_{\text{func}}$ showed weak length-prediction performance. These quantitative results are consistent with the PCA visualization of the latent spaces, where $\mathbf{Z}_{\text{func}}$ separates sequences by class label, $\mathbf{Z}_{\text{length}}$ organizes sequences by length, and $\mathbf{Z}_{\text{res}}$ does not show clear grouping by either attribute (Figure S2). We further quantified target-related information using the mutual-information estimators implemented in scikit-learn. The mutual information between each latent subspace and the target attributes on the held-out test set showed the same pattern: $\mathbf{Z}_{\text{func}}$ had the highest mean mutual information with the binary class label, whereas $\mathbf{Z}_{\text{length}}$ had the highest mean mutual information with peptide length (Table S8). Overall, these results show that PLUM stores class- and length-related information mainly in the intended latent components, while the residual sequence subspace contains less direct class or length information.

Table S8: **Mutual information analysis of PLUM latent subspaces on the held-out test set.** For each latent subspace, mutual information was estimated independently between each latent dimension and the target attribute, and the resulting values were averaged across dimensions. Thus, for an eight-dimensional latent subspace, the reported value represents the mean of the eight dimension-wise mutual information estimates rather than the joint mutual information between the complete latent vector and the target. Binary class-label mutual information was estimated using a nearest-neighbor estimator for discrete targets, whereas peptide-length mutual information was estimated using a nearest-neighbor estimator for continuous targets. Higher values indicate that individual dimensions within the latent subspace contain more recoverable information about the corresponding attribute.

| Latent subspace | Mean MI with binary class label | Mean MI with length |
|---|---|---|
| $\mathbf{Z}_{\text{res}}$ | 0.1026 | 0.0916 |
| $\mathbf{Z}_{\text{func}}$ | 0.6899 | 0.2189 |
| $\mathbf{Z}_{\text{length}}$ | 0.1278 | 2.5840 |
| $\mathbf{Z}_{\text{res}} + \mathbf{Z}_{\text{func}} + \mathbf{Z}_{\text{length}}$ | 0.3068 | 0.9635 |

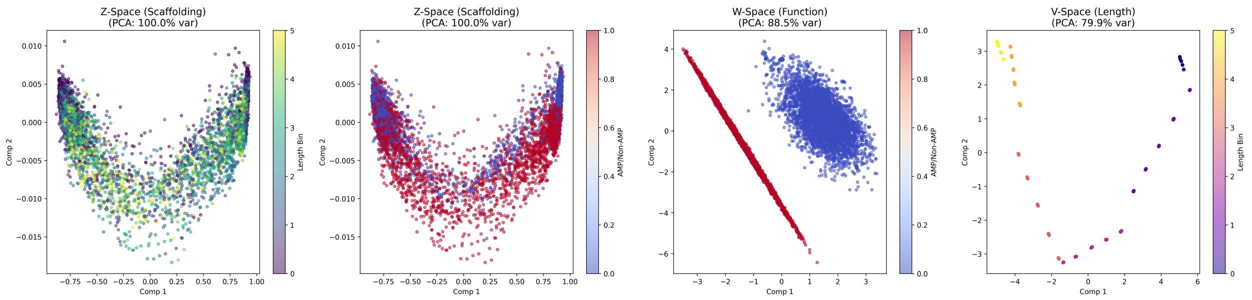


Figure S2: **Structured latent-space organization learned by PLUM.** The trained PLUM encoder splits peptide sequences into three distinct latent spaces to separate different structural traits: sequence variations ($Z$), function ($W$), and sequence length ($V$). For visualization, the high-dimensional positions from each space were compressed to two dimensions using Principal Component Analysis (PCA). (A) The $Z$ **space** is colored by both length and AMP class to demonstrate that general sequence variation remains mixed and unbiased by functional labels. (B) The $W$ **space** is colored by functional class, showing a clear separation between AMP and non-AMP sequences. (C) The $V$ **space** is colored by length bins, highlighting a smooth gradient based on peptide size. Together, these panels show that PLUM successfully organizes and uncouples independent biological properties within its architecture.

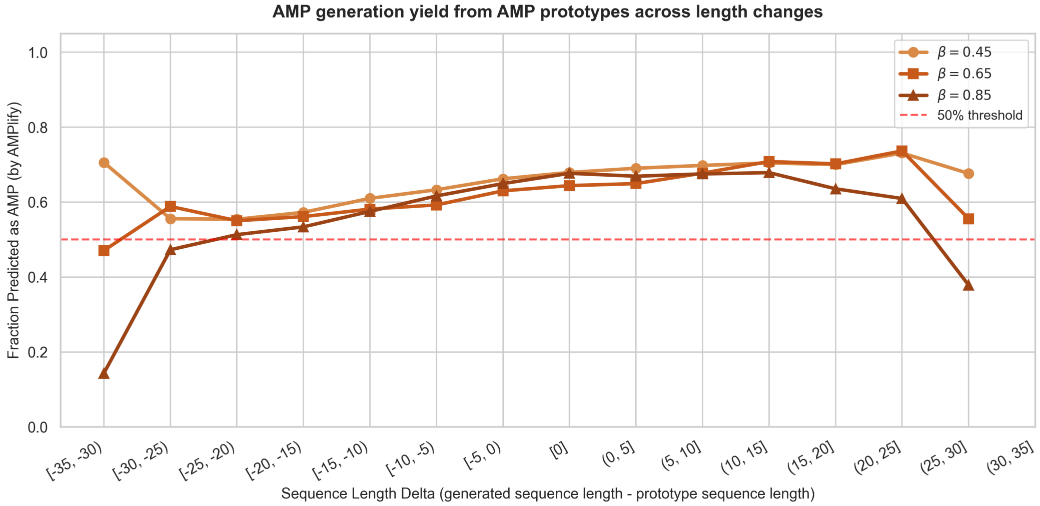


(a) AMP yield from AMP prototypes.

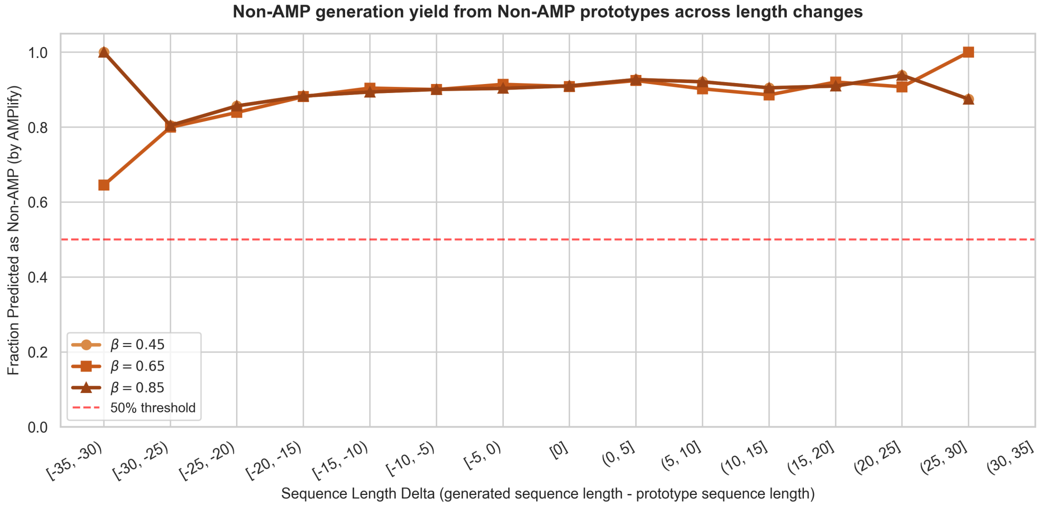


(b) Non-AMP yield from non-AMP prototypes.

Figure S3: **Effect of prototype-to-generated length changes on target-class yield.** In this experiment, the prototype function was kept fixed while PLUM generated peptides at different target lengths relative to the starting prototype. Length delta was defined as the generated sequence length minus the prototype sequence length and grouped into binned ranges. Panel (A) shows AMP generation from AMP prototypes, while panel (B) shows non-AMP generation from non-AMP prototypes. Target-class yield was defined as the fraction of generated peptides classified as the intended class by AMPlify. Curves correspond to three prototype anchoring strengths ($\beta = 0.45$, 0.65, and 0.85), and the dashed red line indicates the 50% threshold.

#### S3.4 Comparative Characterization of Generated Peptides Across Sequence and Physicochemical Properties

The following analyses were conducted on the set of 45,000 generated AMPs and non-AMPs produced by the respective models, as summarized in Table 1 of the main text. These figures provide a detailed evaluation of sequence composition, physicochemical properties, and predicted enzymatic stability. We also include physicochemical properties of the generated peptides from the prototype-guided peptide generation, as shown in Figure 3 from the main text.

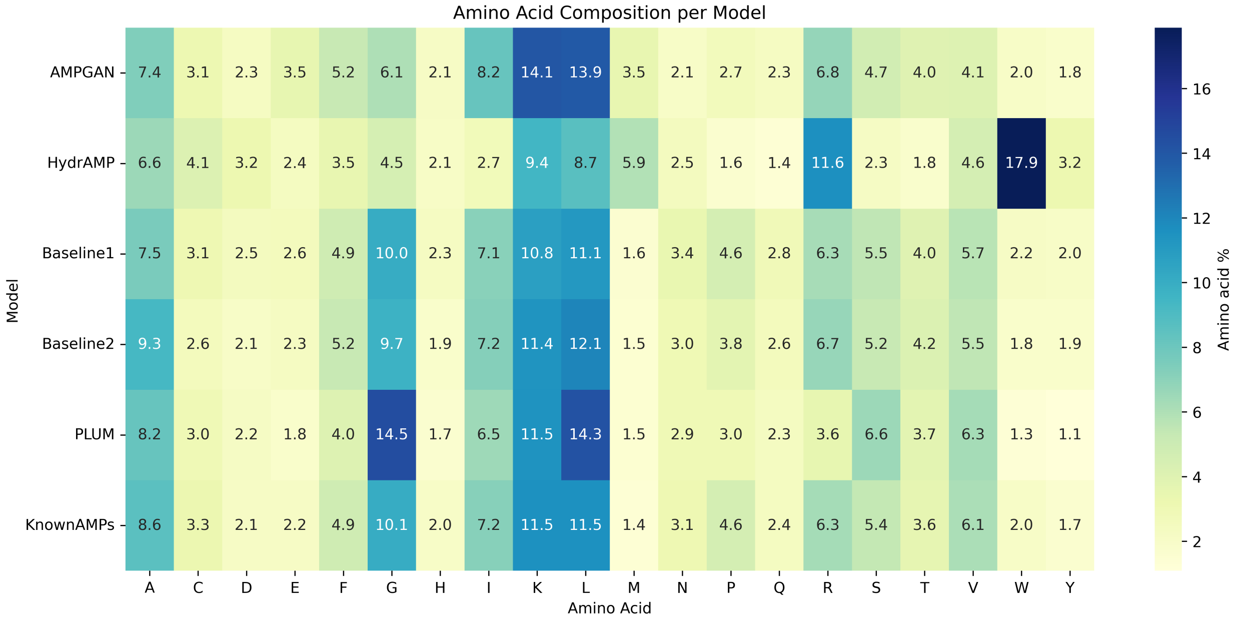


Figure S4: **Amino acid composition** of 45,000 generated AMPs from PLUM, AMPGAN, HydrAMP, Baseline1, and Baseline2 compared with 4,723 AMPs from the reference AMP dataset (KnownAMPs).

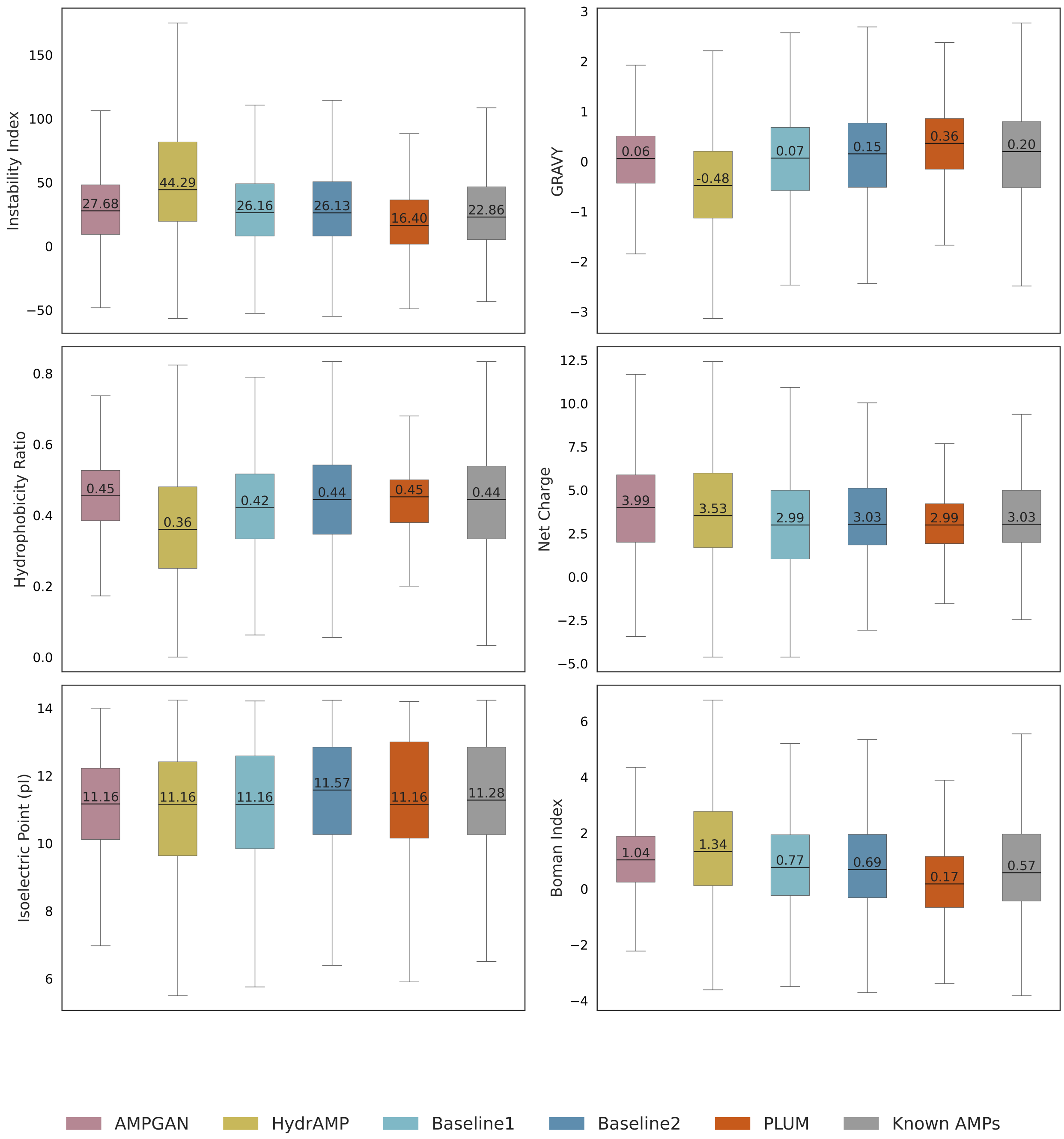


Figure S5: **Comparison of physicochemical property distributions of AMPs.** Distributions of the instability index, GRAVY (Grand Average of Hydropathy), hydrophobic ratio, net charge, isoelectric point (pI), and Boman index were calculated using the `modlamp` python module. The analysis compares 4,723 AMPs from the reference AMP dataset (KnownAMPs) against the 45,000 AMP sequences generated by PLUM, AMPGAN, HydrAMP, Baseline1, and Baseline2. Boxplots depict the median, interquartile range, and overall spread of each property, with median values annotated above each box.

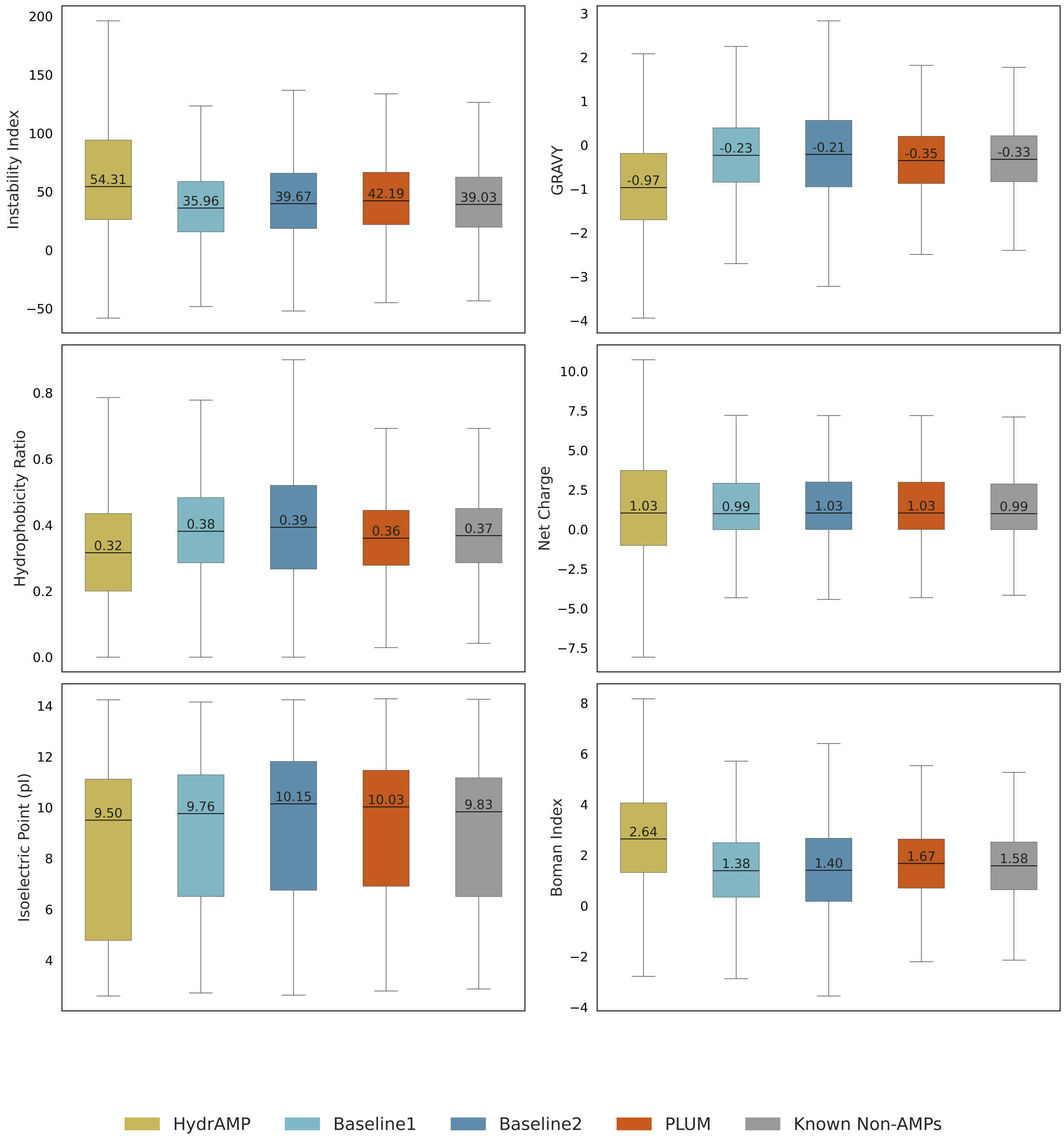


Figure S6: **Comparison of physicochemical property distributions of non-AMPs.** Distributions of the instability index, GRAVY (Grand Average of Hydropathy), hydrophobic ratio, net charge, isoelectric point (pI), and Boman index were calculated using the `modlamp` python module. The analysis compares 4,554 non-AMPs from the reference AMP dataset (KnownNonAMPs) against the 45,000 non-AMP sequences generated by PLUM, AMPGAN, HydrAMP, Baseline1, and Baseline2. Boxplots depict the median, interquartile range, and overall spread of each property, with median values annotated above each box.

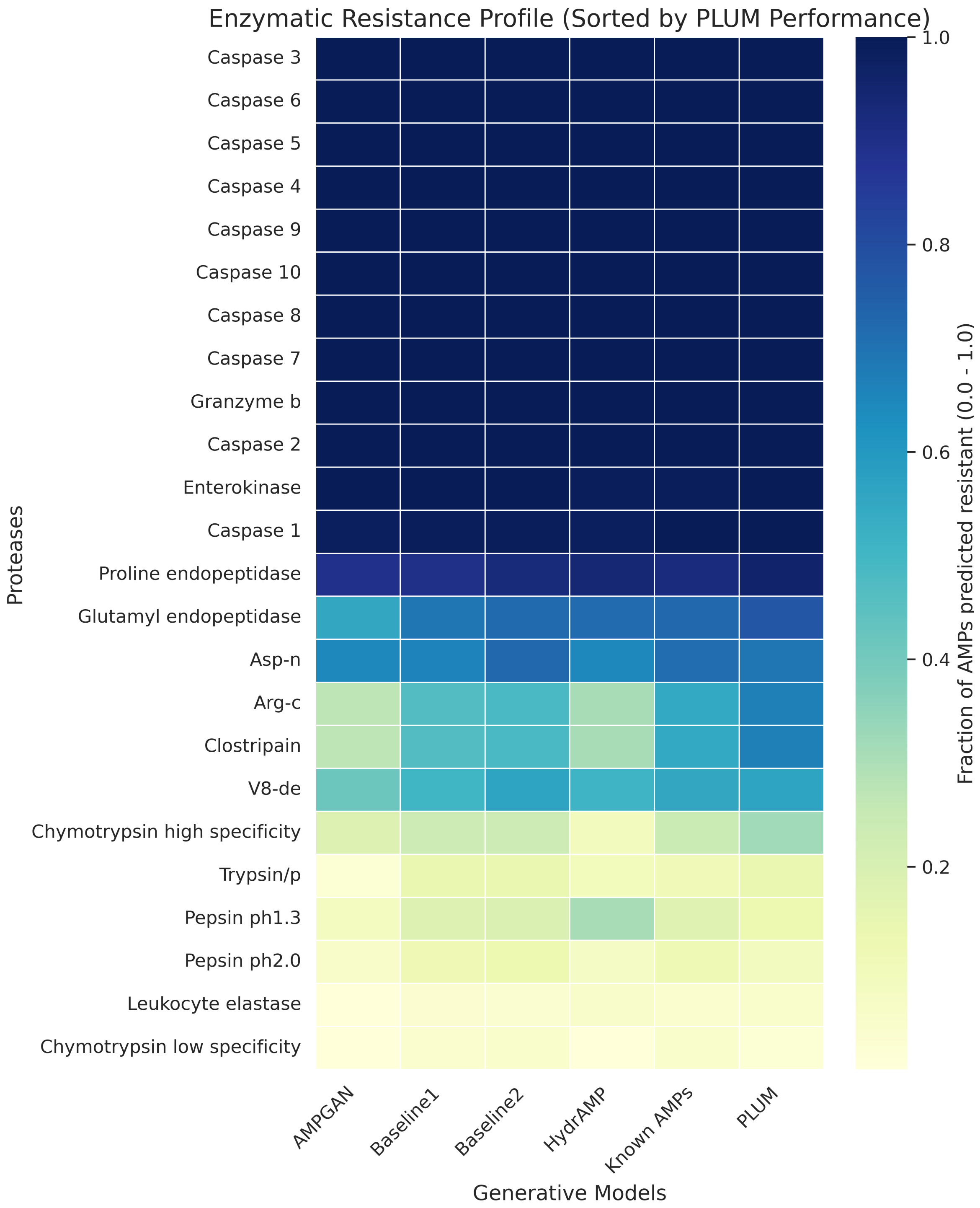


Figure S7: **Predicted proteolytic resistance profile** of 45000 generated AMPs and 4,723 AMPs from the reference AMP dataset (KnownAMPs). Resistance was evaluated across 24 distinct proteases using the Python module `pyteomics`. The heatmap displays the mean resistance fraction for each model, defined as the proportion of sequences resistant to specific protease cleavage sites. Values range from 0.0 (light yellow, complete susceptibility) to 1.0 (deep blue, complete resistance). Proteases are ordered by descending resistance performance in PLUM to highlight stability gains relative to the baseline architectures.

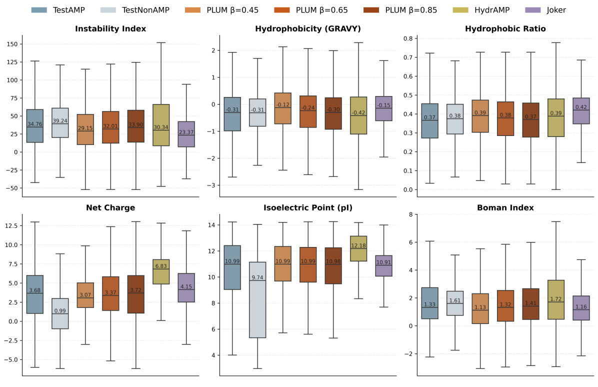


(a) AMP generation from AMP prototypes.

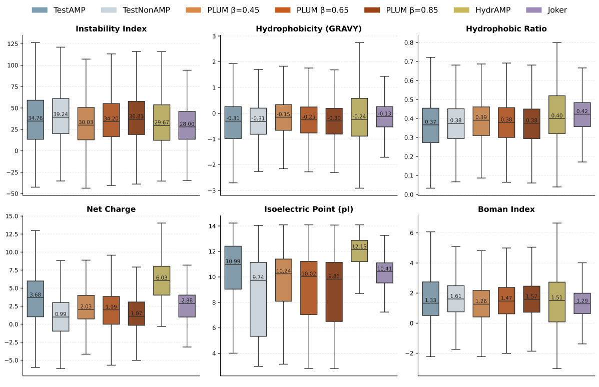


(b) AMP generation from non-AMP prototypes.

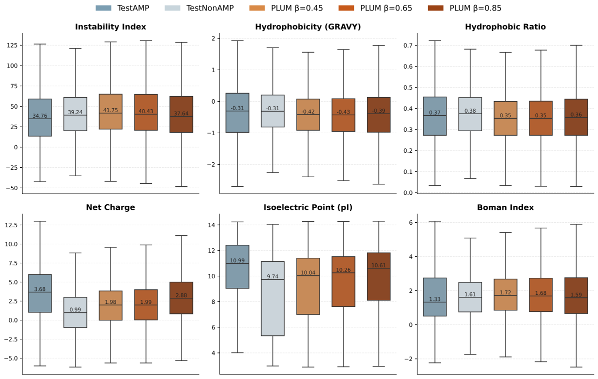


(c) Non-AMP generation from AMP prototypes.

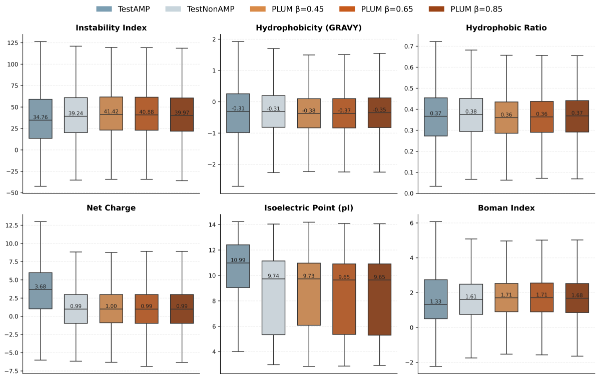


(d) Non-AMP generation from non-AMP prototypes.

Figure S8: Physicochemical characterization of prototype-guided peptide generation across generation tasks and prototype classes.

#### S3.5 Analysis of Generated Sequence Novelty and Internal Diversity

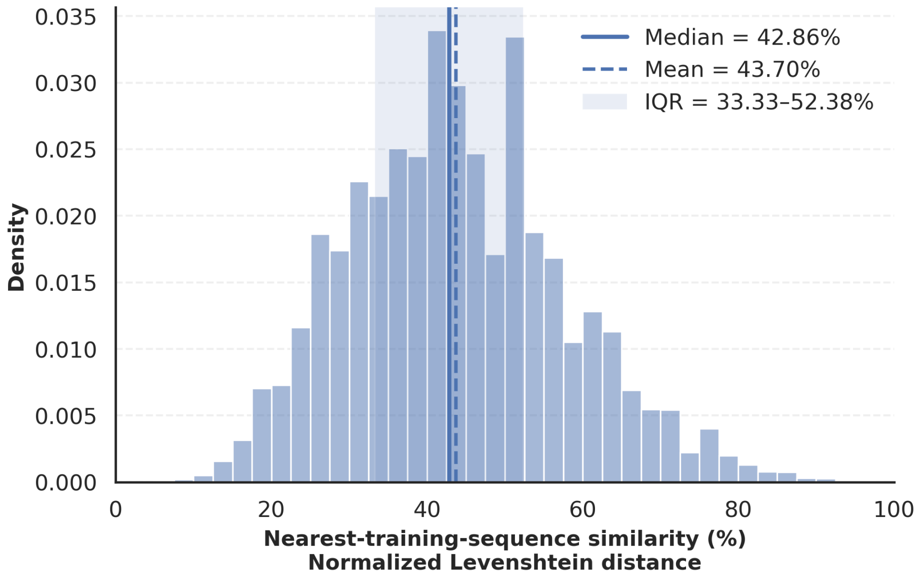


(a) Distribution of normalized Levenshtein similarity between each PLUM-generated AMP and its MMseqs2-selected nearest training-set sequence.

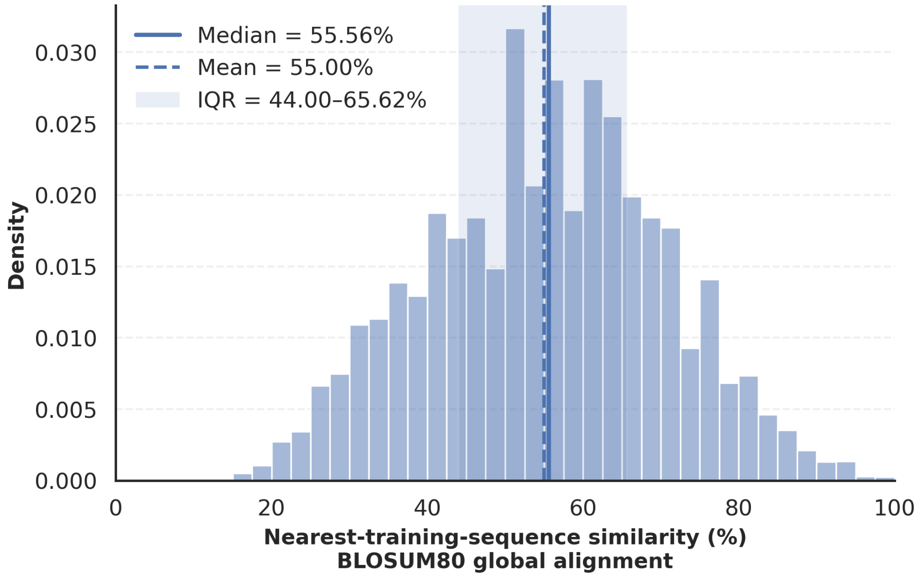


(b) Distribution of BLOSUM80-based amino-acid similarity between each PLUM-generated AMP and its MMseqs2-selected nearest training-set sequence.

Figure S9: **Sequence novelty of PLUM-generated peptides relative to the training set.** Each generated peptide was searched against the positive training-set sequences using MMseqs2, and the highest-scoring candidate training neighbor was selected primarily by bit score, with E-value used as a secondary criterion. The same generated–training pair was then evaluated using two complementary similarity measures. **(A)** Normalized Levenshtein similarity, calculated as $[1 - d_{\mathrm{Lev}}/\max(L_{\mathrm{gen}}, L_{\mathrm{train}})] \times 100$, where $d_{\mathrm{Lev}}$ is the Levenshtein edit distance. **(B)** BLOSUM80 positive-substitution similarity calculated from a global alignment using the BLOSUM80 substitution matrix with gap-opening and gap-extension penalties of $-10$ and $-0.5$, respectively. BLOSUM80 similarity was defined as the percentage of positions in the complete global alignment containing an amino-acid pair with a positive BLOSUM80 substitution score; gap positions contributed to the alignment length but not to the number of similar positions. Vertical lines indicate the mean and median similarity, and the shaded region denotes the interquartile range.

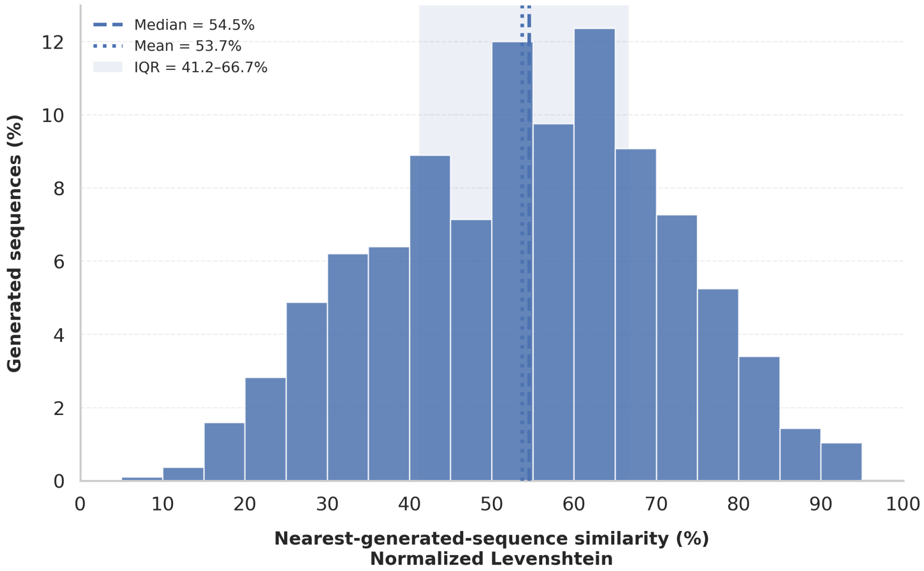


(a) Distribution of normalized Levenshtein similarity between each PLUM-generated AMP and its MMseqs2-selected nearest non-self-generated neighbor.

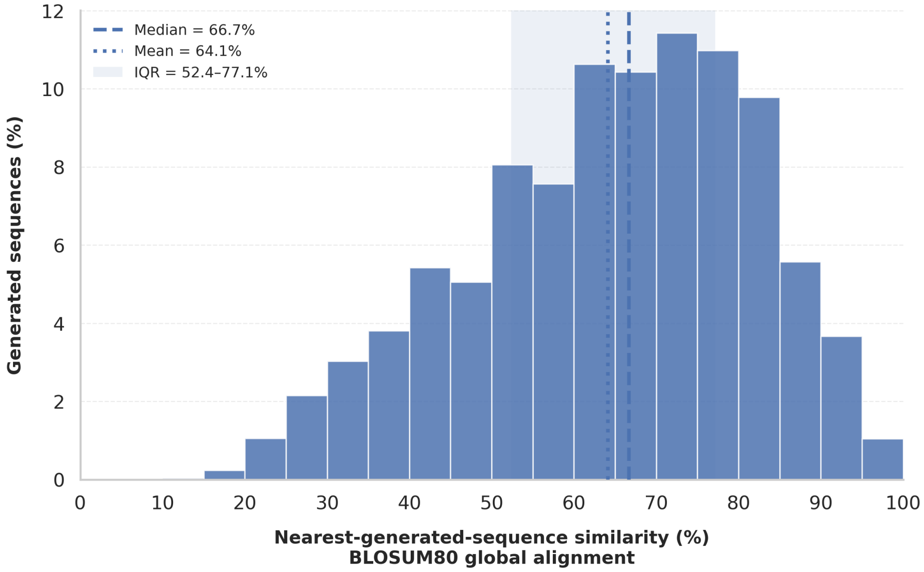


(b) Distribution of BLOSUM80-based amino-acid similarity between each PLUM-generated AMP and its MMseqs2-selected nearest non-self generated neighbor.

Figure S10: **Internal sequence diversity of PLUM-generated AMPs.** The generated peptide set was searched against itself using MMseqs2. Self-record comparisons were removed, and the highest-scoring remaining generated sequence was selected as the nearest non-self neighbor for each peptide. Identical sequences represented by different generated records were retained as valid neighbors. The same generated–generated pair was subsequently evaluated using two complementary similarity measures. **(A)** Normalized Levenshtein similarity, calculated as $[1 - d_{\mathrm{Lev}}/\max(L_1, L_2)] \times 100$, where $d_{\mathrm{Lev}}$ is the Levenshtein edit distance. **(B)** BLOSUM80-based amino-acid similarity calculated from a global alignment using the BLOSUM80 substitution matrix with gap-opening and gap-extension penalties of $-10$ and $-0.5$, respectively. BLOSUM80 similarity was defined as the percentage of positions in the complete global alignment containing an amino-acid pair with a positive BLOSUM80 substitution score, thereby counting both identical residues and conservative amino-acid substitutions as similar. Gap positions contributed to the alignment length but were not counted as similar. Vertical lines indicate the mean and median similarity, and the shaded region denotes the interquartile range. Lower nearest-neighbor similarity indicates greater internal sequence diversity.

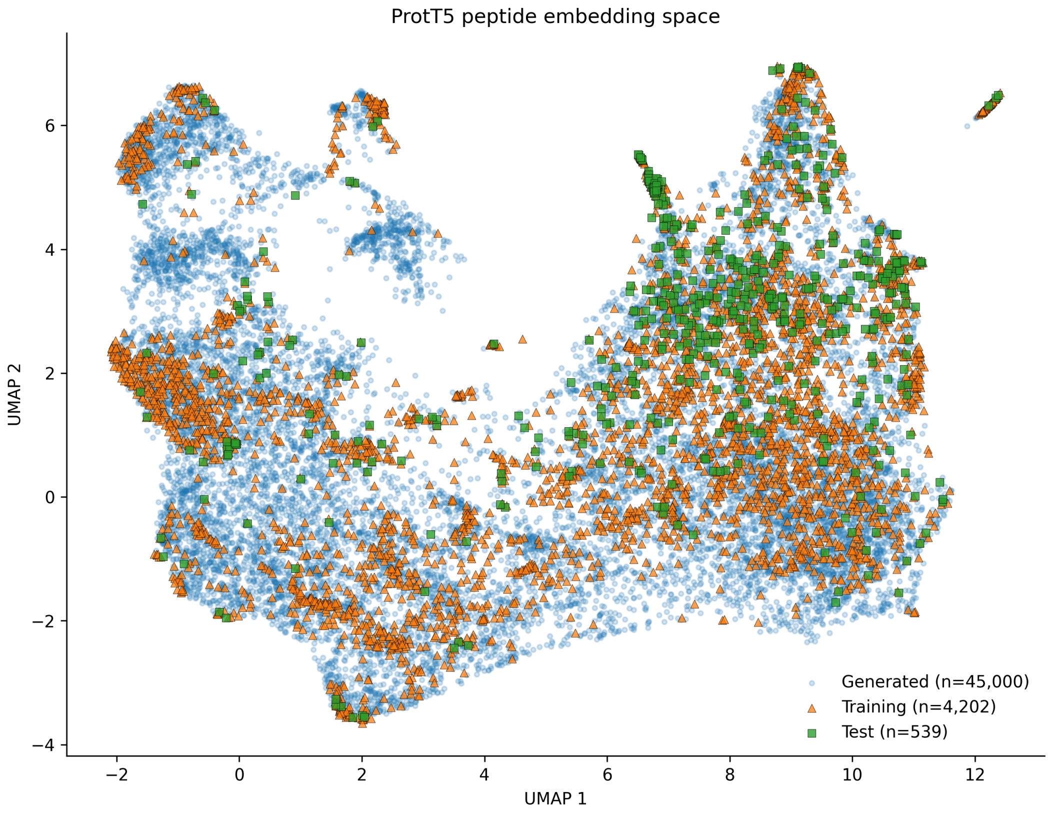


Figure S11: **ProtT5 embedding-space distribution of AMP training, AMP test, and PLUM-generated AMPs.** Peptide sequences from the positive training set, held-out positive test set, and PLUM-generated set were represented using ProtT5 sequence embeddings. Residue-level embeddings were mean-pooled to obtain a single representation for each peptide, followed by dimensionality reduction using principal component analysis (PCA) and two-dimensional UMAP with cosine distance. All sequences were included when constructing the embedding space. For visualization, the complete training and test sets were shown, while up to 15,000 generated peptides were randomly sampled to reduce point-density and visual overlap. Different markers distinguish training, test, and generated peptides. The distribution illustrates the relationship between generated AMPs and experimentally derived AMPs in ProtT5 representation space.

[3] Ulka Gawde, Shuvechha Chakraborty, Faiza Hanif Waghu, Ram Shankar Barai, Ashlesha Khanderkar, Rishikesh Indraguru, Tanmay Shirsat, and Susan Idicula-Thomas. Campr4: a database of natural and synthetic antimicrobial peptides. *Nucleic Acids Research*, 51(D1):D377–D383, 2023.

[4] Tianyue Ma, Yanchao Liu, Bingxin Yu, Xin Sun, Huiyuan Yao, Chen Hao, Jianhui Li, Maryam Nawaz, Xun Jiang, Xingzhen Lao, et al. Dramp 4.0: an open-access data repository dedicated to the clinical translation of antimicrobial peptides. *Nucleic Acids Research*, 53(D1):D403–D410, 2025.

[5] Mohamed F Mohamed, Ahmed Abdelkhalek, and Mohamed N Seleem. Evaluation of short synthetic antimicrobial peptides for treatment of drug-resistant and intracellular staphylococcus aureus. *Scientific reports*, 6(1):29707, 2016.

[6] Nelson G. Oliveira Júnior, Camila M. Souza, Danieli F. Buccini, Marlon H. Cardoso, and Octávio L. Franco. Antimicrobial peptides: Structure, functions and translational applications. *Nature Reviews Microbiology*, 23(11):687–700, 2025.

[7] Malak Pirtskhalava, Anthony A Amstrong, Maia Grigolava, Mindia Chubinidze, Evgenia Alimbarashvili, Boris Vishnepolsky, Andrei Gabrielian, Alex Rosenthal, Darrell E Hurt, and Michael Tartakovsky. Dbaasp v3: database of antimicrobial/cytotoxic activity and structure of peptides as a resource for development of new therapeutics. *Nucleic acids research*, 49(D1):D288–D297, 2021.

[8] Mohammad Rahnamaeian and Andreas Vilcinskas. Short antimicrobial peptides as cosmetic ingredients to deter dermatological pathogens. *Applied microbiology and biotechnology*, 99(21):8847–8855, 2015.

[9] Johan Svenson, Natalia Molchanova, and Christina I. Schroeder. Antimicrobial peptide mimics for clinical use: Does size matter? *Frontiers in Immunology*, 13:915368, 2022.

[10] Christina Wang, Sam Garlick, and Mire Zloh. Deep learning for novel antimicrobial peptide design. *Biomolecules*, 11(3), 2021.

[11] Jacob Witten and Zack Witten. Deep learning regression model for antimicrobial peptide design. *BioRxiv*, page 692681, 2019.

[12] Zhenyu Yang, Xiaoxi Zeng, Yi Zhao, and Runsheng Chen. Alphafold2 and its applications in the fields of biology and medicine. *Signal Transduction and Targeted Therapy*, 8(1):115, 2023.

[13] Chao Zhong, Yongtao He, Jing Zou, Luyang Gao, Jiahui Wang, Jingyi Zhu, Wenjing Xue, Sanhu Gou, Yun Zhang, Hui Liu, et al. An antimicrobial peptide as a potential therapy

for bacterial pneumonia that alleviates antimicrobial resistance. *Nature Communications*, 16(1):10488, 2025.
